# High-resolution whole-genome analysis of sister-chromatid cohesion

**DOI:** 10.1101/2019.12.17.879379

**Authors:** Elena Espinosa, Evelyne Paly, François-Xavier Barre

## Abstract

Sister-chromatid cohesion describes the orderly association of newly-replicated DNA molecules behind replication forks. It plays an essential role in the maintenance and the faithful transmission of genetic information. It is created by DNA topological links and proteinaceous bridges, whose formation and deposition could be potentially affected by many different DNA-binding proteins. However, a mean to analyse local variations in the duration of cohesion on a whole genome was lacking. Here, we present a High-throughput methodology to monitor Sister Chromatid Contacts (Hi-SC2), and show that it permits to analyse locus-specific variations in sister-chromatid cohesion over the whole length of chromosomes.

## Introduction

During replication, sister DNA copies are orderly associated by the formation of DNA topological links and proteinaceous bridges. The phenomenon is referred to as sister-chromatid cohesion. Cohesion facilitates homology search for recombinational repair and ensures that sister chromatids are suitably positioned for segregation (Kleckner et al., 2014; Morales and Losada, 2018). Fluorescence microscopy observations of bulk DNA and of a limited number of genomic positions in a few cellular models suggested that the duration of cohesion varied according to genomic position (Espeli et al., 2008; Fisher et al., 2013; Javer et al., 2013; Kleckner et al., 2004). However, the amount of time necessary to label a given genomic position to observe its replication and segregation dynamics using live fluorescence microscopy and the limited spatial resolution of fluorescence imaging prevents whole-genome high-resolution analyses of the relative cohesiveness of sister DNA copies.

Here, we present a High-throughput methodology to monitor Sister Chromatid Contacts (Hi-SC2), which can be used to analyse locus-specific variations in sister-chromatid cohesion over the whole length of chromosomes at a high resolution. As a proof of principle, we applied it to *Vibrio cholerae*, the agent of the deadly human disease cholera, which is closely related to *Escherichia coli* in the phylogenetic tree of life, the only organism in which cohesion was so far extensively investigated (Bates and Kleckner, 2005; Joshi et al., 2013; Stouf et al., 2013). *V. cholerae* carries homologues of most of the chromosome replication, maintenance and segregation machineries of *E. coli* (Heidelberg et al., 2000). However, its genome is divided on two circular chromosomes, Chr1 and Chr2, whereas *E. coli* harbours a single chromosome (Trucksis et al., 1998; Yamaichi et al., 1999). Chr1 derives from the mono-chromosomal ancestor of *E. coli* and *V. cholerae* whereas Chr2 derives from a domesticated mega-plasmid. Nevertheless, the organisation and positioning of Chr2 participate to the cell cycle of the bacterium (Galli et al., 2016). Chr1 and Chr2 harbour a single origin of bi-directional replication, *ori1* and *ori2*, respectively. Replication terminates in a region opposite of their origin, next to a site dedicated to the resolution of topological problems due to their circularity, *dif1* and *dif2*, respectively (Galli et al., 2019; Val et al., 2008). Most strains of *V. cholerae* are not pathogenic or only cause local outbreaks of gastroenteritis. Epidemic strains have so far been found in only 2 out of >200 possible somatic O-antigens serogroups (Chatterjee and Chaudhuri, 2003). Epidemic strains are characterized by the capacity to produce cholera toxin, which is responsible for the deadly diarrhoea associated with the disease, and Type IV toxin-coregulated pili (TCP), which are essential for small intestine colonization (Herrington et al., 1988). Both virulence factors are acquired by horizontal gene transfer. Cholera toxin is encoded in the genome of a lysogenic phage, CTXΦ (Waldor and Mekalanos, 1996), which almost always integrates behind a Toxin Linked Cryptic satellite phage, TLCΦ, at the *dif1* locus (Hassan et al., 2010; Midonet et al., 2019). TCP is encoded in a 39.5 kbp vibrio pathogenicity island inserted in Chr1, VPI-1. In addition to its role in intestine colonization, TCP serves as a receptor for CTXΦ (Davis and Waldor, 2003). Hi-SC2 revealed that cohesion is highly prolonged in large genomic territories centred on the origin and terminus of replication of Chr1 and Chr2. It further revealed that the H-NS nucleoid-structuring protein, which acts as a transcriptional repressor in horizontally-acquired elements (Dorman, 2007), dramatically extends the duration of cohesion within VPI-1 and the O-antigen region (Ayala et al., 2015; Chun et al., 2009; Herrington et al., 1988). Finally, Hi-SC2 unveiled small locus-specific variations in the duration of cohesion over the whole length of each of the two *Vibrio cholerae* chromosomes, which had so far escaped scrutiny.

## Results

### Hi-SC2 design

Hi-SC2 exploits paired-end deep-sequencing to determine in parallel the status and position of sister-chromatid contact (SC2) reporters inserted at random positions in a library of cells (Figure 1A). The SC2 reporters are site-specific recombination cassettes. They consist of a very short DNA segment flanked by two directly-repeated site-specific recombination (SSR) sites. The small distance between the SSR sites prevents intramolecular recombination, which would excise the DNA segment separating the SSR sites and leave a single SSR site as a scar. However, intermolecular recombination events between sister copies of a reporter can also lead to the replacement of a cassette by a single SSR site (Demarre et al., 2014; Lesterlin et al., 2012). Hence, the frequency of replacement of a reporter by a single SSR site, ***f_SC_*_2_**, reflects the frequency of contacts between sister chromatids at its position of insertion, which can be used as a measure of the duration of cohesion (Figure 1A). In brief, Hi-SC2 starts with the introduction of a conditional expression allele of site-specific recombinase in a cell line and the creation of a library of cells harbouring a cognate SC2 reporter at random genomic positions by transposition. Production of the SSR is induced for different lengths of time during cell growth and/or at specific stages of the cell cycle. High-throughput paired-end sequencing is then used to determine the position and recombination status of the SC2 reporters.

**Figure 1.**
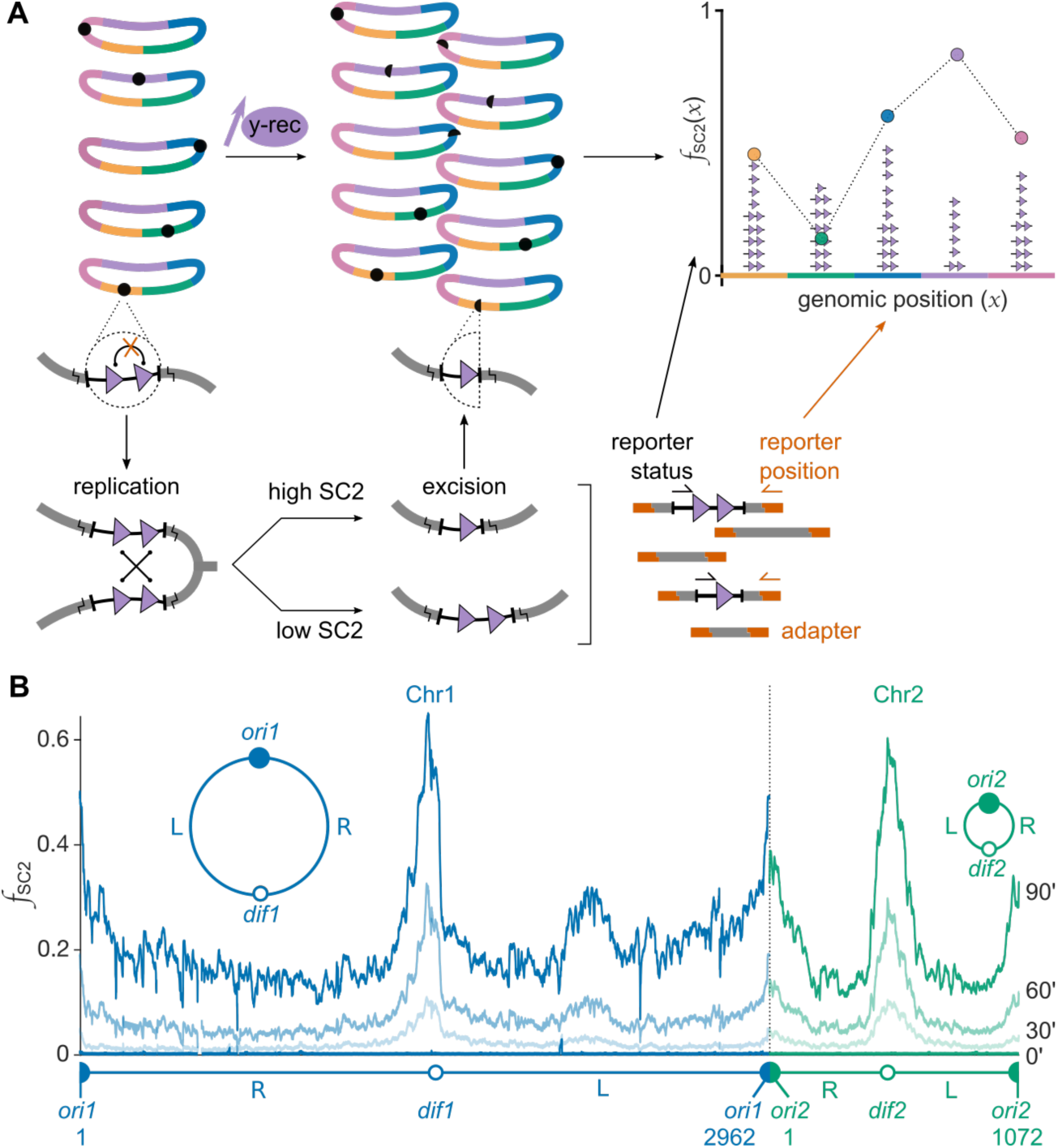
Overview of Hi-SC2. (A) Schematic representation of the assay: the SC2 reporter (black circle) consists of two directly repeated site-specific recombination (SSR) sites (purple triangles), separated by a very short piece of DNA. When the cognate tyrosine recombinase is produced (y-rec), recombination events between sister chromatids drive the replacement of the two SSR sites at the SC2 reporter position by a single one. High-throughput paired-end sequencing permits to determine the genomic position of the SC2 reporter (orange primer) and the number of SSR sites at this position (black primer). The proportion of single SSR sites to the total number of SC2 reporter inserted at a given genomic position, ***x***, reflects the frequency of sister-chromatid contacts at this position, ***f_SC_*_2_**(***x***). (B) Cre-based ***f_SC_*_2_** along *V. cholerae* Chr1 (blue lines) and Chr2 (green lines). Solid and open circles depict the origin and terminus of replication of the chromosome, respectively; the left and right replication arms are indicated as L and R, respectively. Chromosome sizes are indicated in kbp.

### Analysis of Sister-Chromatid Cohesion in *V. cholerae*

As a proof of principle, we applied Hi-SC2 to *V. cholerae*. To this end, we equipped an El Tor N16961 *V. cholerae* strain with an inducible allele of the tyrosine recombinase of phage P1, Cre, which acts on a 34 bp SSR site, *loxP*. We inserted a SC2 reporter consisting of two directly repeated *loxP* sites separated by 21 bp in a Mariner transposon specifically designed for efficient Tn-seq analysis (van Opijnen et al., 2009). We created cell libraries containing >10^5^ independent transposition events, which corresponds to an average of >1 reporter every 40 bp (Table S1). A total number of 10^7^ paired-end sequencing reads was sufficient to obtain a mean number of >100 reads per reporter position, which permitted to calculate ***f_SC_*_2_** at most (if not all) reporter positions (Figure S1). ***f_SC_*_2_** can be analysed at different scales of precision (Figure S2). For the purpose of the present work, we calculated the mean of ***f_SC_*_2_** over 10 kbp sliding-windows, which permits to highlight both short-range and long-range variations in cohesion along the two *V. cholerae* chromosomes in a small figure (Figure S2). By convention, we present the ***f_SC_*_2_** profile of each chromosome in a clockwise orientation starting from their origin of replication (Figure 1B).

Before Cre production, we didn’t observe any replacement of the SC2 reporter cassette by a single SSR site, demonstrating that such events exclusively resulted from Cre-mediated recombination (Figure 1B, 0’). ***f_SC_*_2_** increased as a function of time after the induction of Cre production (Figure 1B). It was not constant along the length of the genome, reaching a maximum of ∼70% after 90’ in slow growth conditions (Figure 1B). Higher frequencies were obtained at the same time in fast growth conditions, as expected because of the higher number of replication cycles (Figure S3A and B). Independent Cre-based SC2 reporter libraries yielded identical ***f_SC_*_2_** profiles, demonstrating the reproducibility of the technique (Figure S3C and D). By comparison, the excision of a long *loxP*-cassette, which permits Cre-mediated intramolecular recombination, was constant over the whole genome and reached a frequency of 80% in only 30’ of induction in slow growth, demonstrating that ***f_SC_*_2_** profiles were not due to differences in the accessibility of DNA and/or the activity of Cre along the genome (Figure S4).

### Cohesion is extended in large territories centred on *ori1*, *ori2*, *dif1* and *dif2*

***f_SC_*_2_** was strikingly elevated in four large 200 kbp genomic territories centred on the replication origin and replication terminus of each of the two *V. cholerae* chromosomes, indicating that sister copies of these regions remained cohesive for a longer period of time than sister copies of the rest of the chromosome replication arms (Figure 1, Figure S3 and S5). ***f_SC_*_2_** was maximal near *dif1* and *dif2*, (Figure 1B, Figure S3 and S5). It reached up to 70% at these positions after 90’ of induction in slow growth conditions, while it was only in the order of 20% in the flanking regions (Figure 1B, Figure S3 and S5). The extensive cohesion of the *dif1*-terrirory fits with previous fluorescence microscopy observations, which suggested that sister *dif1* copies remained associated until the initiation of septum constriction (David et al., 2014; Demarre et al., 2014; Galli et al., 2017). However, sister copies of *dif2*, which is replicated at the same time of the cell cycle as *dif1*, were found to split ahead of the initiation of septum constriction, in apparent contradiction with the Hi-SC2 data (Demarre et al., 2014; Galli et al., 2019). To resolve this issue, we analysed sister-chromatid contacts at the time of cell division using a reporter based on a site-specific recombination machinery whose activity is restricted to the time of constriction, Xer (Demarre et al., 2014; Kennedy et al., 2008; Val et al., 2008). Xer-based ***f_SC_*_2_** was null at all genomic positions but in the immediate vicinity of *dif1* and *dif2*, in agreement with the ordered choreography of segregation of Chr1 and Chr2 and the temporal restriction of the activity of Xer (Figure 2). It remained very low over the whole *dif2*-territory, including the *dif2* position (Figure 2). It was also low in most of the *dif1*-territory, apart from the immediate proximity of *dif1*, where it was as high as Cre-based ***f_SC_*_2_** (Figure 2). Taken together, these results suggest that sister copies of the origin and terminus regions of the two *V. cholerae* chromosomes are highly cohesive behind the replication forks and that most of the links between sister copies of genomic positions within the *dif1-* and *dif2*-territories are released prior to the initiation of cell division, with the exception of the immediate vicinity of *dif1* (Figure 2).

**Figure 2.**
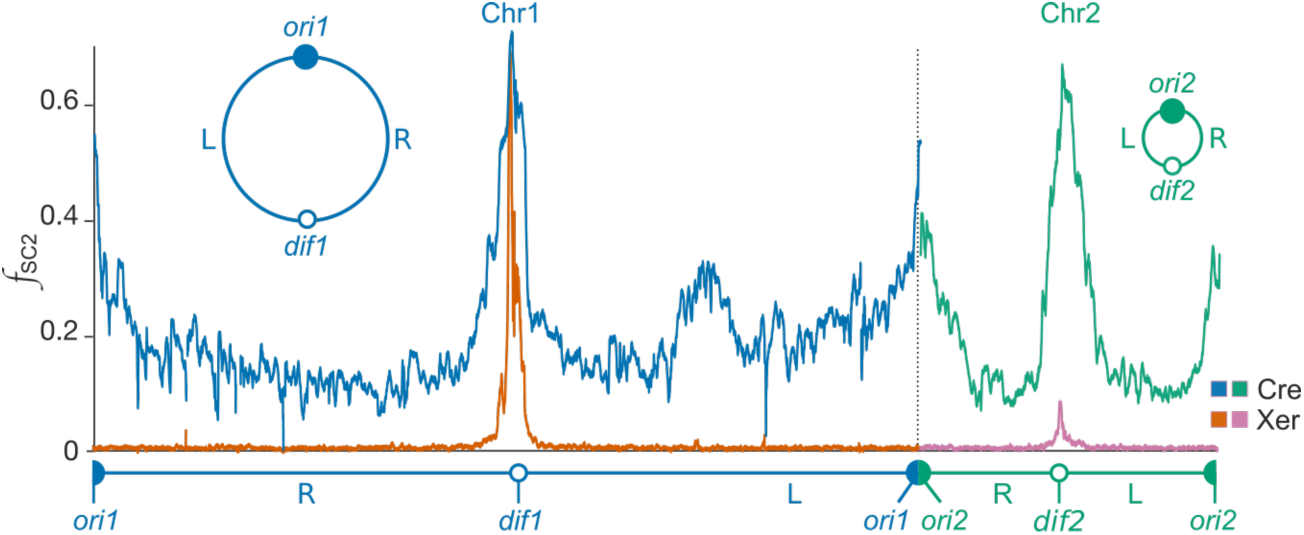
Relative cohesiveness of sister chromatids during cell division. Comparison of the Cre-based (FX224 dataset) and Xer-based (FX129 dataset) Hi-SC2 profiles. Cre permits to monitor ***f_SC_*_2_** from the time of replication of a genomic position to the time of separation of the resulting sister copies. Xer permits to specifically monitor ***f_SC_*_2_** during the final stage of cell division, septum constriction.

### VPI-1 creates a highly cohesive territory

Hi-SC2 further revealed the existence of a large 200 kbp territory in the middle of the left replication arm of Chr1 where ***f_SC_*_2_** was elevated (Figure 3A and Figure S3D). This domain is centred on VPI-1. Deletion of VPI-1 supressed the elevated ***f_SC_*_2_** within the whole territory, without affecting ***f_SC_*_2_** in the rest of the genome (Figure 3A and Figure S6). These results prompted us to use fluorescence microscopy to inspect the segregation of a locus within the VPI-1 territory, L1, with respect to the segregation of a locus immediately downstream of it on the same replication arm, L2 (Figure 3B). As a comparison, we analysed the relative segregation of two genomic positions in the right replication arm, R1 and R2, which are located at approximately the same distance from *ori1* than L1 and L2, but had equivalent ***f_SC_*_2_** values (Figure 3B). We observed more cells with two R1 foci and a single R2 focus than cells with a single R1 focus and two R2 foci, as expected from the earlier replication of R1 (Figure 3B, left panel). In contrast, there were more cells with a single L1 focus and two L2 foci than cells with two L1 foci and a single L2 focus, demonstrating that the separation of sister L1 copies was delayed compared to the separation of sister L2 copies, as predicted by Hi-SC2 (Figure 3B, middle panel). Furthermore, we found that deletion of VPI-1 alleviated the segregation delay of sister L1 copies (Figure 3B, right panel). Together, these results demonstrate that VPI-1 creates a highly cohesive territory.

**Figure 3.**
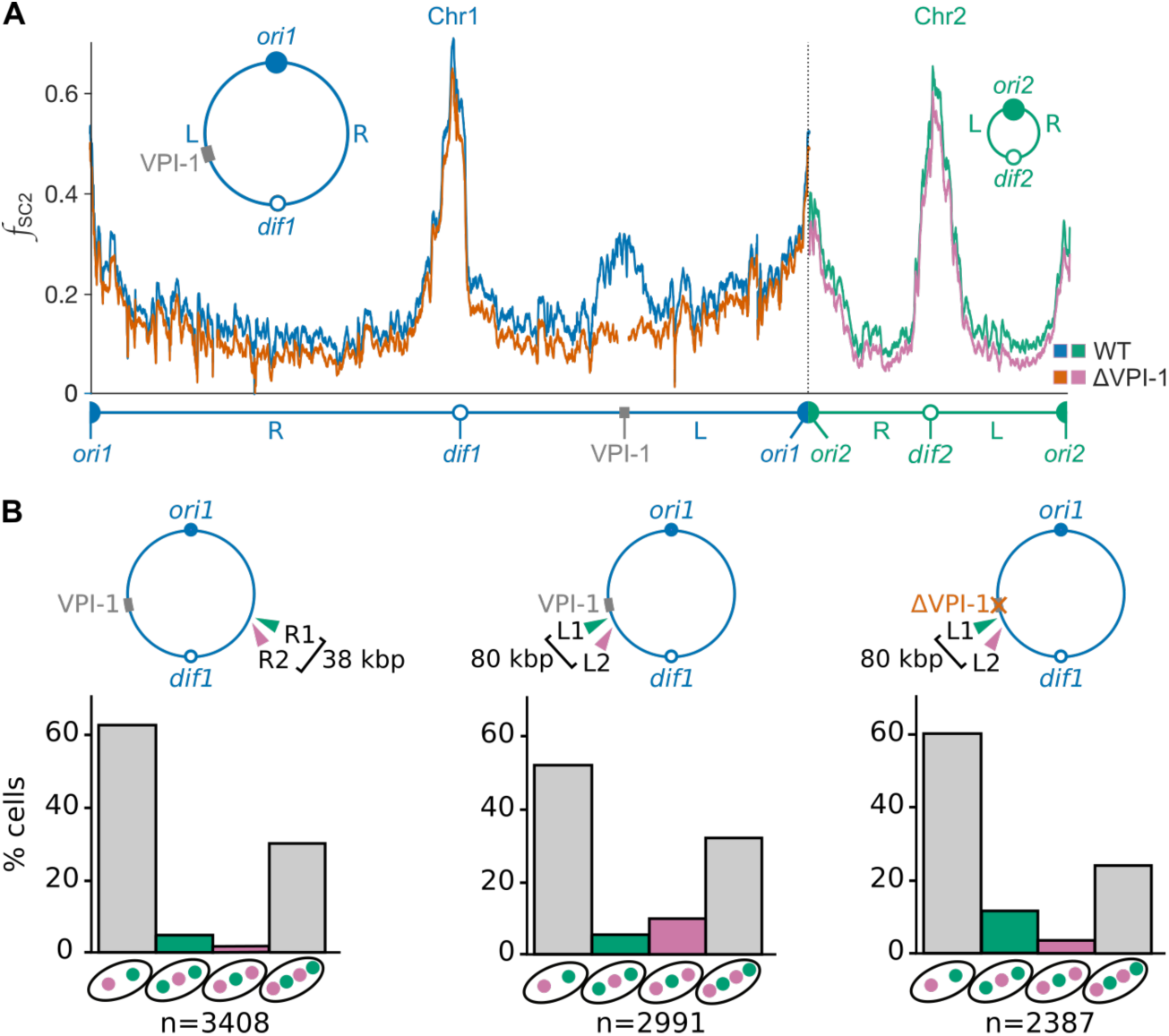
A highly cohesive territory surrounds the VPI-1 Vibrio pathogenicity island. (A) Comparison of the Cre-based Hi-SC2 profiles in WT and ΔVPI-1 cells. The position of VPI-1 is indicated with a grey box (from bp 2176653 to bp 2217966 kbp). (B) Fluorescence microscopy analysis of the order of segregation of two neighbouring loci on the same replication arm. Origin- and terminus-proximal loci are depicted in green and purple, respectively. Graphs show the relative proportion of cells with the indicated number of foci. Informative cell categories are highlighted in colour. n: total number of analysed cells from two independent experiments. Distances between markers are indicated.

### The Histone-like Nucleoid Structuring protein promotes cohesion

The Histone-like Nucleoid Structuring protein, H-NS, acts as a transcriptional repressor in horizontally-acquired elements (Dorman, 2007). It regulates the virulence of *V. cholerae* and covers the VPI-1 and O-antigen regions (Figure 4, bottom panel; Ayala et al., 2015; Chun et al., 2009; Herrington et al., 1988). Deletion of *hns* suppressed sister-chromatid cohesion within the 100 kbp origin-proximal part of the VPI-1 territory and the O-antigen region without affecting surrounding regions, demonstrating that H-NS is a region-specific cohesion factor (Figure 4 and Figure S7).

**Figure 4.**
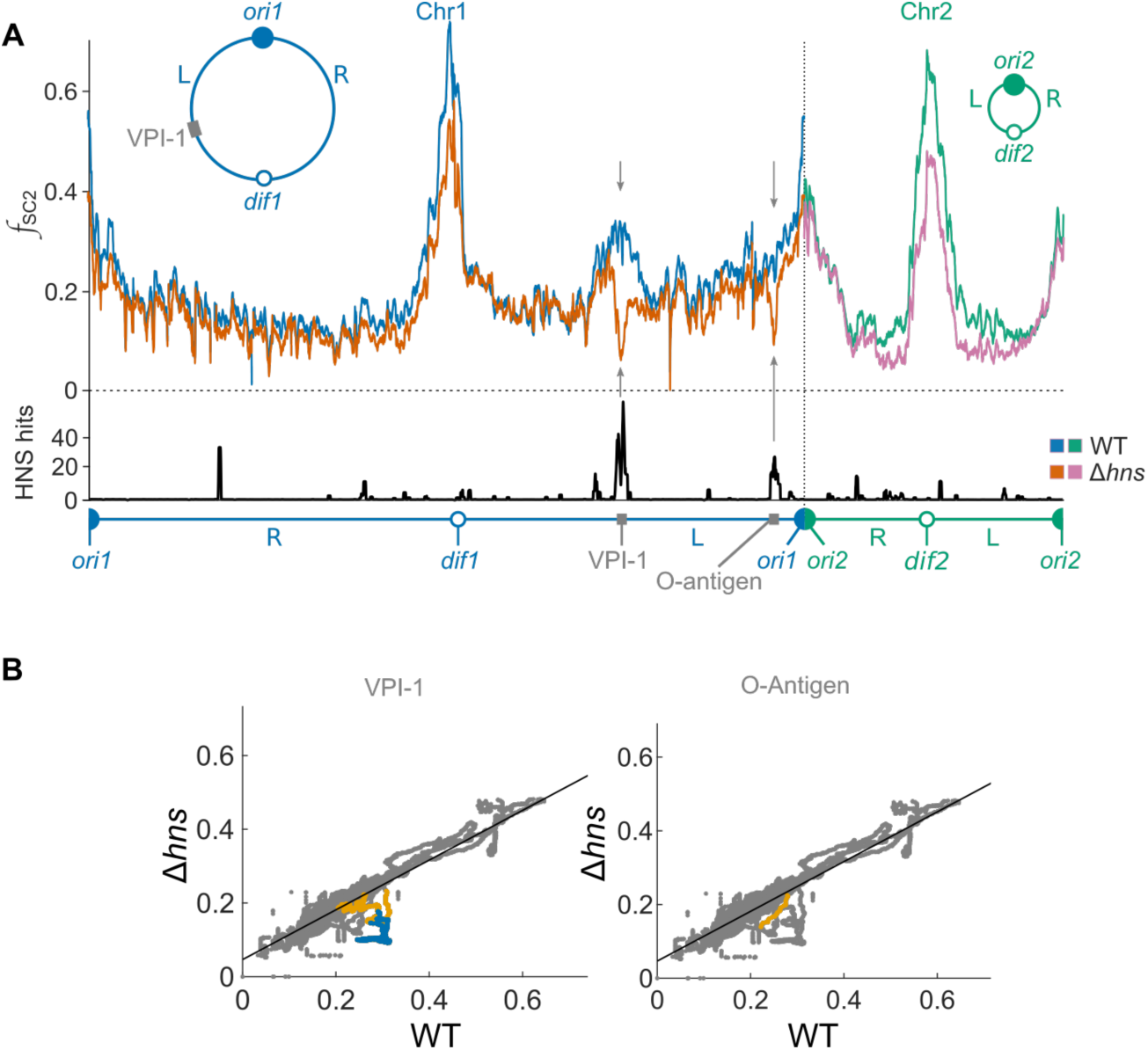
H-NS extends the cohesiveness of sister chromatids within VPI-1 and the O-antigen region. (A) Upper panel: Cre-based Hi-SC2 profiles in WT and Δ*hns* cells; Lower panel: relative distribution of H-NS along the *V. cholerae* genome analysed by ChIPseq (Ayala et al., 2015). Grey arrows indicate regions of significantly lower SC2 frequencies in Δ*hns* cells. (B) Correlation between the SC2-reporter cassette excision frequencies in Δ*hns* (FX215 dataset) and WT cells (FX224 dataset). Black line: linear regression of the Δ*hns* and WT data. Left panel: data points within VPI-1 and the 2160 to 2283 kbp origin-proximal part of the VPI-1 territory are highlighted in blue and orange, respectively; Right panel: data points within the 2825477 to 2839408 bp O-antigen region are highlighted in orange.

### Hi-SC2 unveiled small locus-specific variations in the duration of cohesion

Small short-distance (<10 kbp) fluctuations in the Hi-SC2 profiles are superimposed on the large long-distance ***f_SC_*_2_** variations (Figure 1B, Figure S2). Independent Cre-based SC2 reporter libraries yielded identical short-distance ***f_SC_*_2_** variations, indicating that they were not due to experimental noise (Figure S3C, S6 and S7). Furthermore, short-distance ***f_SC_*_2_** variations changed depending on the growth conditions, indicating that they were linked to physiological parameters (Figure S3A). Taken together, these results suggest that short-distance ***f_SC_*_2_** variations probably reflect small locus-specific differences in the duration of cohesion over the whole length of chromosomes, which couldn’t be explored with current fluorescence microscopy techniques.

## Discussion

Sister-Chromatid Cohesion plays an essential role in the maintenance and the faithful transmission of genetic information. Previous work suggested that many different DNA-binding proteins could affect cohesion. However, the duration of cohesion was only characterized in detail at a limited number of genomic positions in a few model organisms. The Hi-SC2 methodology we describe here offers the possibility to analyse local variations in the duration of cohesion at a high resolution over a whole genome (Figure 1). It gives a massive amount of information, but data analysis and representation are relatively simple. The methodology can be used to monitor the relative amount of time during which sister DNA copies remain cohesive during the entire cell cycle (Figure 1) or to analyse cohesion at specific stages of the cell cycle by placing the SC2 recombinase under an inducible promoter or choosing it for its specific cell-cycle regulation (Figure 2). Its application to *V. cholerae* demonstrated that cohesion is connected to other genome management processes, such as replication (Figure 2) and transcription regulation (Figure 3 and 4).

### Initiation and termination of replication take place in cohesive territories

Hi-SC2 demonstrated that sister-chromatid cohesion is prolonged in 200kbp territories centred on the replication origin and replication terminus of each of the two *V. cholerae* chromosomes (Figure 2). Thus, Hi-SC2 opens up a new field of investigation on the relationship between the formation of these highly cohesive territories, the actual replication process and the regulation of replication in different bacterial and eukaryotic models.

### Cohesion is released before the actual repositioning of sister copies

Fluorescence microscopy showed that sister copies of the origin region of the *E. coli* chromosome are highly cohesive (Bates and Kleckner, 2005; Joshi et al., 2013). We could have expected cohesion at the origin region of the *V. cholerae* chromosomes to be more limited than cohesion at the origin of the *E. coli* chromosome because the *V. cholerae* chromosomes, as most other bacterial chromosomes and in contrast to the *E. coli* chromosome, are equipped with a partition machinery dedicated to the active segregation of sister origin copies (Fogel and Waldor, 2006; Yamaichi et al., 2007). The observation of the elevated cohesiveness of the replication origins of the *V. cholerae* chromosomes suggest that the partition machineries cannot release all of the cohesion links but only serve to assure the correct repositioning of sister origins in the cell after their individualisation.

Sister copies of the terminus region of the chromosome of slow growing *E. coli* cells were also proposed to be highly cohesive based on genetic experiments and fluorescence microscopy observations (Galli et al., 2017; Stouf et al., 2013). It was attributed to the requirement for the activity of a conserved ATP-dependent DNA pump anchored in the division septum, FtsK, in sister termini separation (Stouf et al., 2013). However, FtsK is only active during septum closure (Demarre et al., 2014; Kennedy et al., 2008), whereas our Hi-SC2 results demonstrate that sister copies of the terminus region of Chr2 and of the major part of the terminus region of Chr1 are separated before the initiation of constriction (Figure 2). This observation suggests that FtsK does not participate in the release of cohesion at the terminus of the *V. cholerae* chromosomes but only serve to assure their faithful distribution in opposite daughter cell compartments.

Taken together, these results suggest that segregation is a multi-step process, with release of cohesion and condensation driving the individualisation of sister copies before their active repositioning in the cell, as proposed in eukaryotes.

### Transcription regulation programs can promote cohesion

Hi-SC2 revealed that VPI-1 created a highly cohesive territory in the middle of the left replication arm of Chr1 (Figure 3). VPI-1 is a major pathogenic island of *V. cholerae*, which was known to be regulated by H-NS. Deletion of H-NS alleviated the cohesiveness of sister DNA copies within VPI-1 and the origin proximal part of the highly cohesive territory surrounding it (Figure 4). Deletion of H-NS also reduced cohesion in the O-antigen region, even if this region is located in the origin domain of Chr1 (Figure 4). The implication of a bacterial repressor in cohesion creates a parallel with the action of cohesins in eukaryotes, which ensure sister-chromatid cohesion but also participate in the regulation of transcription via the formation of topologically associated domains (Uhlmann, 2008).

### Hi-SC2 unveiled small locus-specific variations in the duration of cohesion

The Hi-SC2 methodology permits to monitor the relative frequency of contacts between sister copies at an unprecedented high resolution (Figure S3). It unveiled short-range locus-specific variations in cohesion, which could not be observed using fluorescence microscopy (Figure 1-4). We are attracted by the idea that small locus-specific ***f_SC_*_2_** variations could be at the origin of the cyclic accumulation and relief of mechanical stress observed during the individualization of sister chromatids (Bates and Kleckner, 2005; Fisher et al., 2013). The Hi-SC2 methodology will permit to investigate the multiple factors at the origin of the local variations in cohesion along the length of chromosomes, including the local transcriptional status for the genomic positions, the proteins binding to them, their composition in bases and their methylation.

## Experimental Procedures

### Bacterial strains and growth conditions

Strains are listed in Extended Data Table 2. All *V. cholerae* strains were derivatives of El Tor N16961 (Heidelberg et al., 2000) and were constructed by natural transformation (Meibom et al., 2005). Plasmids used for the construction of strains are listed in Table S3. Strains were grown in LB at 37° C for fast growth conditions or M9 media supplemented with 0.2% fructose at 30°C for slow growth conditions.

### Cre- and Xer-based SC2 reporters

Cre and Xer are topologically independent recombinases, i.e. they can recombine sites belonging to different molecules, recombine sites directly repeated on the same molecule and sites in inverted orientations. The Cre- and Xer-based SC2 reporters consist of DNA cassettes flanked by two directly-repeated *loxP* or *dif1* sites, respectively. The recombination sites were separated by 21 bp in the case of *loxP* and 27 bp in the case of *dif* to prevent their excision by intramolecular recombination.

### Cre-activity reporter

The Cre-activity reporter consists of a DNA cassette flanked by two directly-repeated *loxP* sites separated by 205 bp. It is more than the persistence length of DNA, which permits excision by intramolecular recombination.

### Tightly inducible Cre and Xer production alleles

To ensure no leaky production of Cre or Xer, the recombinase genes were placed under the control of the arabinose promoter (P_BAD_) and the *E. coli lacZ* promoter was inserted at the end of the gene in inverse orientation (invP_lac_). Transcription from invP_lac_ inhibits leaky transcription from P_BAD_ in the absence of arabinose. IPTG was added to all cultures to ensure invP_lac_ transcription before the time of induction of the recombinases. To further ensure no leaky production, the Cre and Xer production alleles were inserted as a single copy in the *V. cholerae* genome.

### Reporter library construction

The Cre- and Xer-based SC2 reporters and the Cre-activity reporter were randomly inserted by transposition. The transposition plasmids used to insert the Cre- and Xer-based SC2 reporters and the Cre-activity reporter are derivatives of the pSWT23 suicide plasmid conjugation vector (Table S3; (Demarre et al., 2005)). They carry the transposase of the *magellan5* Mariner transposon and a kanamycin resistance marker flanked by the inverted repeats of the transposon (*Himar*) in which a *Mme*I restriction site has been inserted to facilitate the analysis of the position of insertion of the kanamycin cassette by high-throughput sequencing (van Opijnen et al., 2009). The Rd2SP sequence (Illumina) was introduced downstream of the kanamycin resistance for sequencing purposes. The Cre- and Xer-based SC2 reporters and the Cre-activity reporter were cloned downstream of the Rd2SP sequence. The transposition vectors were introduced by conjugation as follows: donor (EE14 for the Cre-based SC2 reporter and Cre-activity reporter; EE6 for the Xer-based SC2 reporter) and recipient cells were grown for 16 hours at 37 °C in LB. The donor was grown with DAP (0.3 mM) and chloramphenicol (25 µg/ml) whereas the recipient was grown with zeocin (25 µg/ml) and IPTG (0.1 mM). 250 µl of each culture were mixed, washed in LB with IPTG, resuspended into 30 µl of LB with IPTG and placed into a 0.45µm filter (Millipore) in a LB agar plate supplemented with DAP and IPTG. Conjugation was carried out for 4 hours at 37°C. Thirty conjugations were performed per each library. After conjugations, cells were pooled together, resuspended in 6 ml of LB with IPTG and plated on LB plates containing kanamycin (50 µg/ml) and IPTG. Plates were incubated for 16 hours at 30 °C except for EEV113 and EEV114, which were incubated for 40 hours. Cells were scraped, mixed together in LB with IPTG and stored in aliquots at −80 °C.

### Sister-Chromatid Cohesion assays

Aliquots of the transposon library were thawed on ice and ∼10^9^ cells were diluted into 100 ml of media supplemented with IPTG. Cells were grown until exponential phase (OD_600_ 0.2). Then, ∼10^9^ cells were centrifuged and transferred into 10 ml of fresh media without IPTG. After 30 minutes, arabinose was added at a final concentration of 0.02%. ∼10^9^ cells were recovered and frozen in liquid nitrogen at different time points after induction. For the essential gene analysis, cells were collected after 24 hours (FX409 and FX410 datasets).

### Paired-end sequencing

Extraction of chromosomal DNA was performed using the Sigma GenElute bacterial genomic DNA kit. 2 µg of DNA were digested for 4 hours with *Mme*I (NEB) following the manufacturer’s instructions. Digested DNA was purified with MiniElute columns (Qiagen). The Pippinprep (SAGE science) was used to purify digested fragments of a length comprised between 800 bp and 2 kbp, which corresponds to the region containing the transposon. 400 ng of purified DNA was ligated overnight at 16 °C to adapters (Table S4) with T4 DNA ligase (NEB). Ligated DNA was purified using MiniElute columns (Qiagen). For each library, 3 PCR reactions were performed using 2 µl of ligated DNA as template and a pair of Illumina oligos (Table S4, P5 and P7). The PCR reactions were performed using the following parameters: 98 °C for 30 seconds, 17 cycles of 98 °C 30 seconds, 68 °C 30 seconds, 72 °C 30 seconds and a final step of 72 °C for 7 minutes. PCR products were concentrated with a MiniElute columns (Qiagen) and purified using Pippinprep (SAGE science). Paired-end sequencing was performed on a NextSeq (Illumina).

### Sequencing data analysis

First, barcodes and adapter sequences were removed from the sequences using Cutadapt version 1.9.1 (Martin, 2011). Second, Cutadapt was used to remove the transposon end sequence. Third, Cutadapt was used to select total reads and reads in which the reporter cassette had been excised. Finally, reads were aligned on the *V. cholerae* EEV29 genome with the BWA software (Li and Durbin, 2010).

### Hi-SC2 analysis

Unless otherwise stated, ***f_SC_*_2_** was calculated as the ratio between the number of single SSR sites over a 10 kbp window around the position and the total number of transposition insertion in this window. Note, however, that ***f_SC_*_2_** can also be analysed at the bp level. Points for which we recovered less than 500 reads, because the 10 kbp window contains too many essential genes or repeated sequences were deleted (Figure S1). Hi-SC2 profiles were calculated using MatLab (Mathworks). Scripts are available on request.

### Essential genes analysis

Essential genes and intergenic regions were determined using ARTIST (Pritchard et al., 2014). Results are shown in Table S6.

### Fluorescence microscopy analysis

To confirm that the elevated frequency of cassette excision within the VPI-1 domain corresponded to a local increase in cohesion, we tagged a locus in the VPI-1 domain, just downstream of the pathogenicity island itself, L1, and a locus outside of the VPI-1 domain, L2, 80 kbp downstream of L1. As a control, we tagged two loci on the right replication arm of Chr1, R1 and R2, at a similar distance from *ori1* than L1 and L2, respectively. Hi-SC2 analysis suggested that the cohesiveness of sister R1 and R2 copies were similar. L1 and R1 were tagged with a *parSPMT1* site. L2 and R2 were tagged with a 3 kbp *lacO* array. The positions and numbers of L1 and R1 foci were visualized with a ParB_PMT1_-YGFP fusion (David et al., 2014; Stouf et al., 2013) and the positions and numbers of L2 and R2 loci were visualized with a LacI-mCherry fusion (Lau et al., 2003). Differences in timing of segregation under slow growth were determined by calculating the proportion of cells displaying 1 YGFP and 1 mCherry foci (1:1), 2 YGFP and 1 mCherry foci (2:1), 1 YGFP and 2 mCherry foci (1:2) and 2 YGFP and 2 mCherry foci (2:2). Cells from the 1:1 and 2:2 categories are not informative. However, the proportion of cells belonging to the 2:1 and 1:2 categories reflect the difference in timing of separation of sister DNA copies at the two loci (Stouf et al., 2013). Cells from the 1:2 or 2:1 category are not very abundant, which imposed the analysis of >1000 cells. Cells were grown in M9 media supplemented with fructose as described in the reporter cassette excision assays. Cells were placed on an agarose pad (1% w/v) (Ultrapure, Invitrogen). For snapshot analysis, cells images were acquired using a DM6000-B (Leica) fluorescence microscope equipped with an ORCA-Flash 4.0 camera (Hamamatsu) and piloted with MetaMorph (Version 7.8.8.0, Molecular Devices). They were analysed using MicrobeTracker (Sliusarenko et al., 2011).

### Data and materials availability

Hi-SC2 Sequence data was deposited in the ArrayExpress Archive of Functional Genomics Data (https://www.ebi.ac.uk/arrayexpress/).

## Acknowledgements

We thank C. Possoz, G. Lelandais, V. Lioy and Y. Yamaichi and the high-throughput sequencing facility of the I2BC. The work was supported by the ERC (FP7/2007-2013, grant number 28159) and the ANR (grants 2016-CE12-0030-0 and 2018-CE12-0012-03).

---

**Figure S1.**
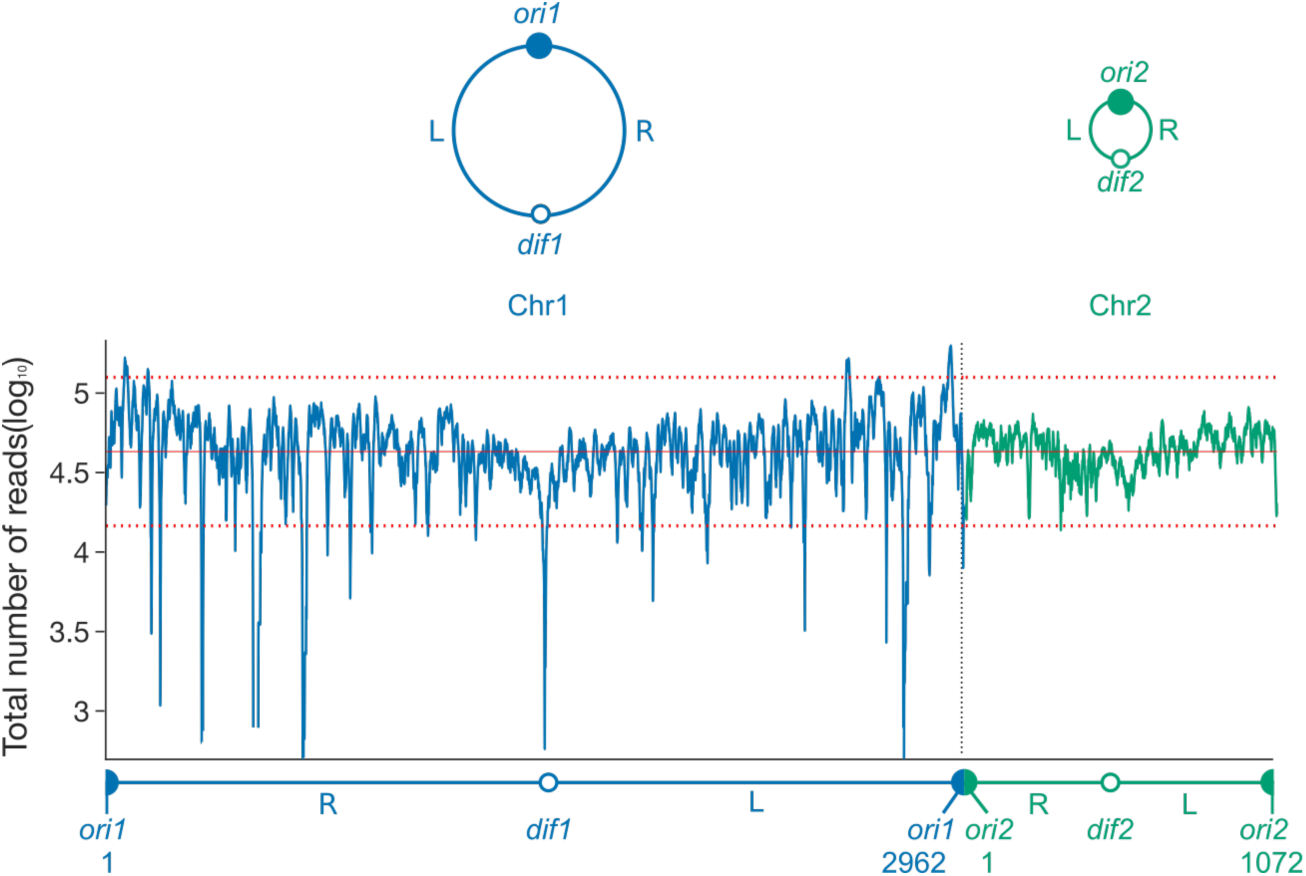
Typical SC2 reporter coverage map. The figure shows the total number of reads of the SC2 reporter in a 10 kbp region surrounding each genomic position at the 90’ time point of the experiment presented in Figure 1B (FX224 dataset). Read numbers are shown in log scale. The few positions where the number of reads was reduced correspond for the most part to repeated sequences, such as the ribosomal genes, which cannot be mapped. Note that >500 reads were obtained in the 10 kbp surrounding almost all genomic positions, which is sufficient to estimate *f*_SC2_ at this resolution. Chr1 and Chr2 schematic maps and Chr1 and Chr2 insertion data profiles are shown in blue and green, respectively. The length of the chromosomes is indicated in kbp on the linear map. The chromosome coordinates are set so that the region of the origin of replication of each chromosome is at the start and end of the linear map representation (filled circles). The region of their terminus of replication region is located in the middle of the linear map (open circles). Plain and doted horizontal lines show the position of the mean (4.64) ± twice the standard deviation (0.23) of the data.

**Figure S2.**
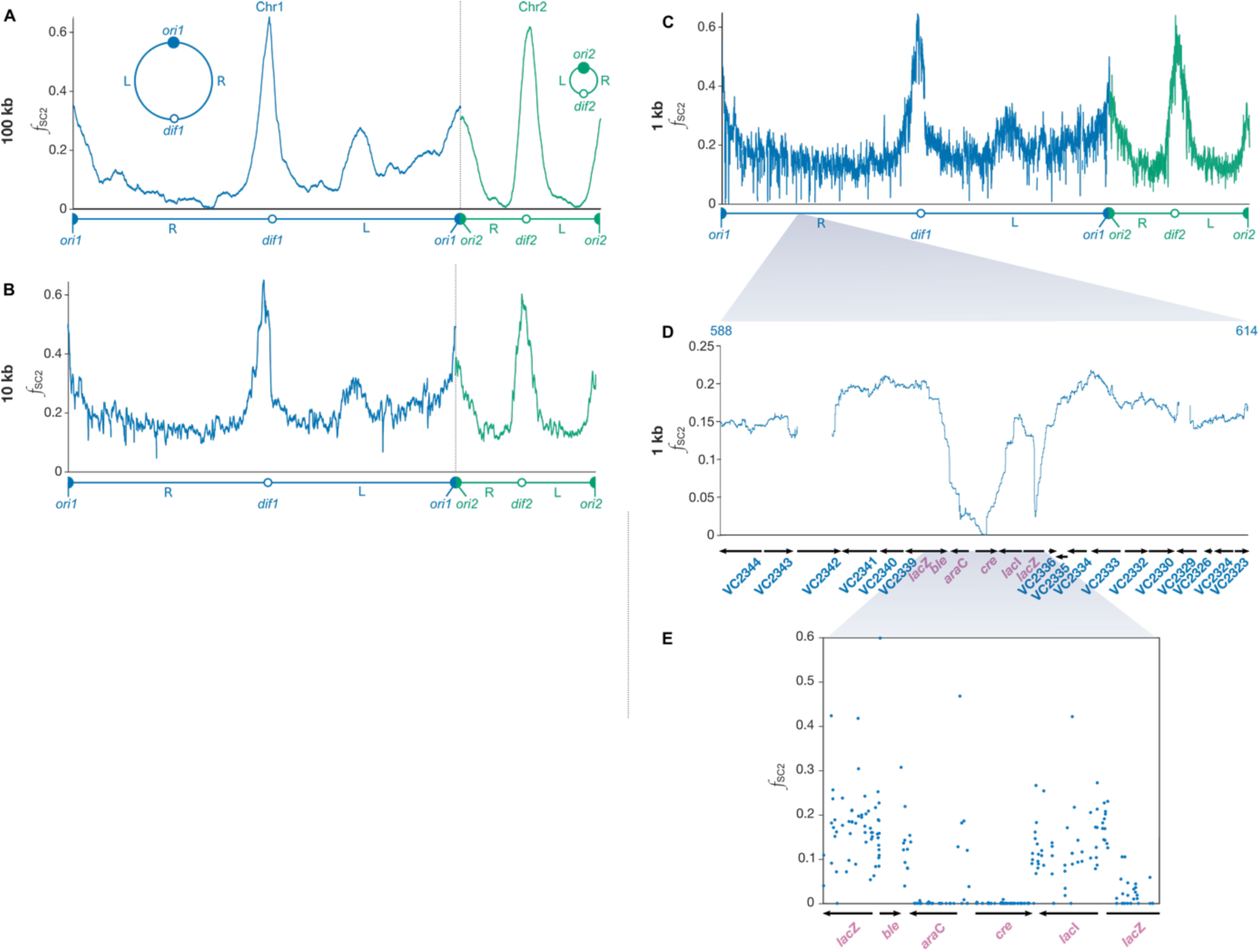
Short-range and large-range variations in sister chromatid cohesion. Hi-SC2 data can be represented using 100 kbp (A), 10 kbp (B) or 1 kbp (C) sliding-windows to highlight large-range and short-range variations in the relative frequency of sister chromatid contacts. The data corresponds to the 90’ time point of the Cre-based assay shown in Figure 1 (FX224 dataset). (D) Zoom of the 1 kbp excision profile around the region of insertion of Cre (from position 588 to 615 kbp). (E) Zoom on the frequency of excision around the region of insertion of Cre (from position 598254 to 604418 bp), showing the frequency of excision at each position of insertion of the SC2 reporter. As expected, no excision is observed for reporters inserted in the *cre* and *ara*C genes.

**Figure S3.**
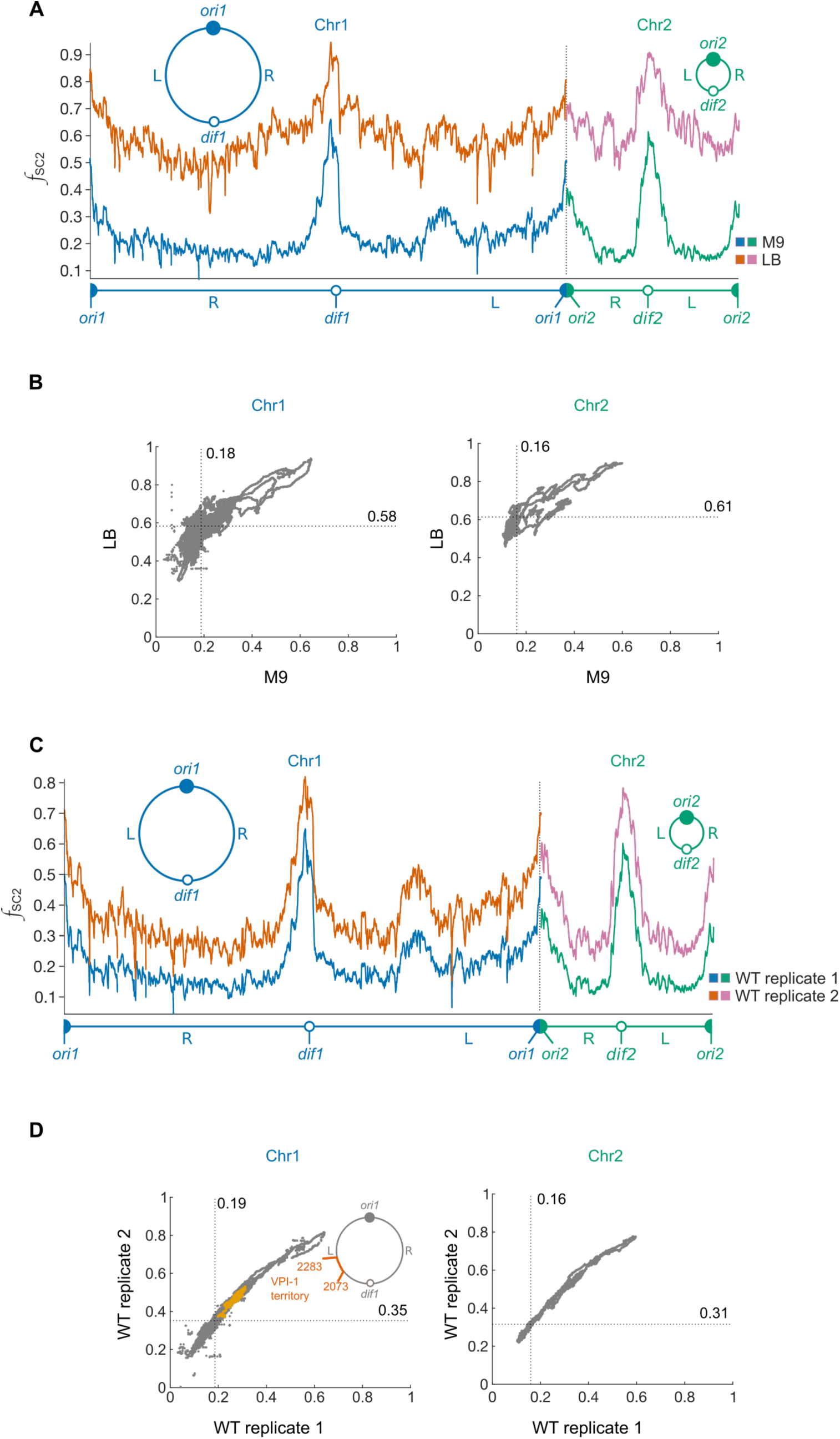
Validity of Hi-SC2 analysis. (A) Comparison of Cre-based *f*_SC2_ in cells under slow growth conditions (M9) and cells under fast growth conditions (LB). The data of the slow growth conditions corresponds to the 90’ time point of Figure 1 (FX224 dataset, blue and green lines). The data of the fast growth conditions was obtained after 60’ of induction of Cre (FX129 dataset, orange and pink lines). Despite the shorter induction time, fast-growth *f*_SC2_ was higher than slow-growth *f*_SC2_ at all genomic positions, as expected from the increase in the number of replication cycles. The difference in the Hi-SC2 profiles under slow- and fast-growth is particularly evident in the middle of the replication arms of the two chromosomes. (B) Cre-based *f*_SC2_ under fast growth versus Cre-based *f*_SC2_ under slow growth. The horizontal and vertical dotted lines indicate the position of the median *f*_SC2_ in the FX129 and FX224 dataset, respectively. (C) Reproducibility of Hi-SC2. Blue and green lines: Hi-SC2 profile obtained under slow growth condition after 90’ of Cre-induction shown in Figure 1 (FX224 dataset); Orange and pink lines: Hi-SC2 profile obtained for a second transposition library under the same experimental conditions (FX239 dataset). The *f*_SC2_ of the FX239 dataset were slightly higher than those of the FX224 dataset, which probably reflects a difference of growth between the two experiments. Note, however, that long-distance variations and short-distance fluctuations were almost identical. (D) Cre-based *f*_SC2_ from the FX239 dataset vs Cre-based *f*_SC2_ from the FX224 dataset. The horizontal and vertical dotted lines indicate the position of the median *f*_SC2_ in the FX239 dataset and the FX224 dataset, respectively. The positions corresponding to the VPI-1 territory are highlighted in orange (from position 2176653 to position 2217966).

**Figure S4.**
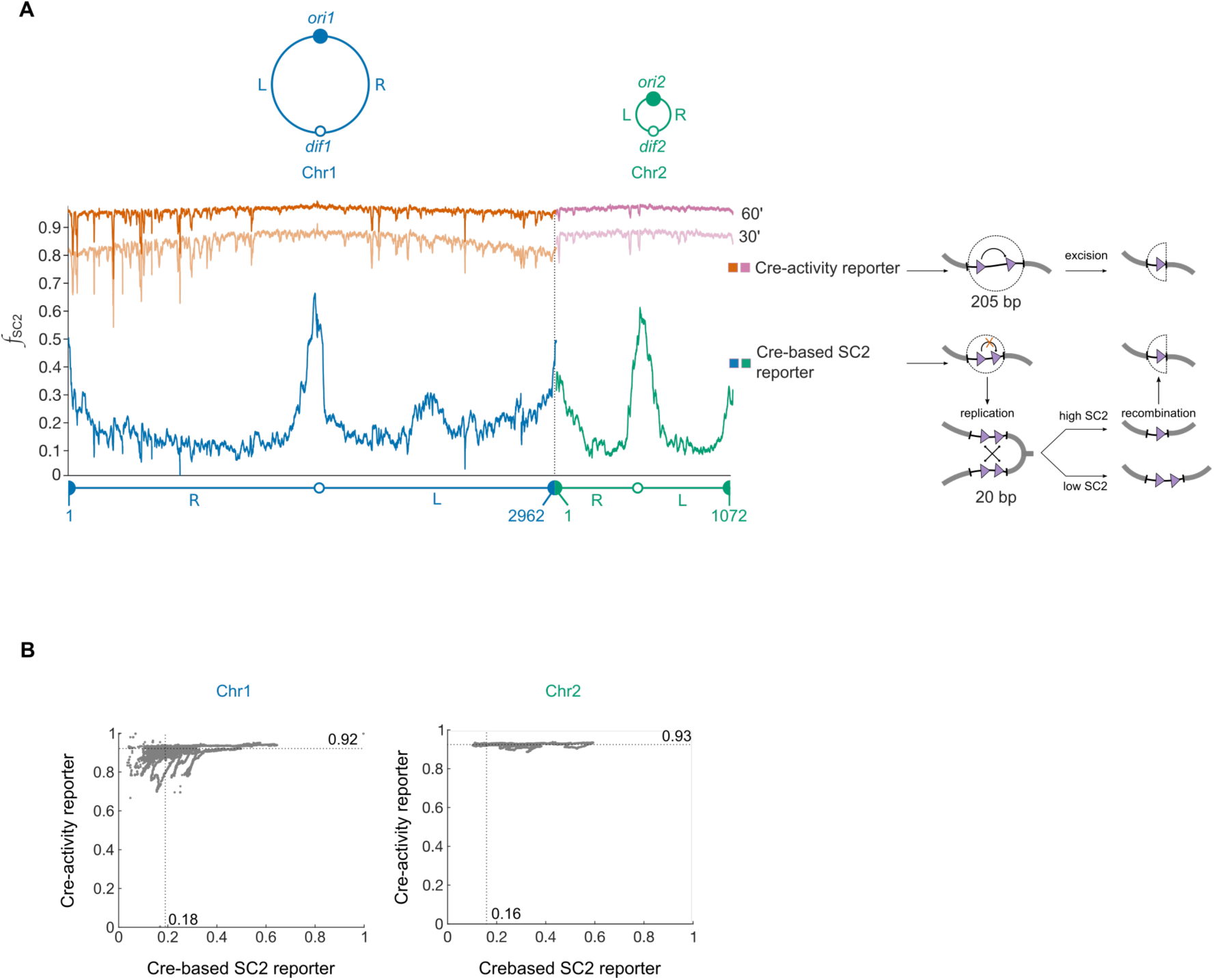
Accessibility of genomic positions to Cre and activity of Cre along the genome. (A) Cre-activity reporter cassette excision frequency profiles obtained under slow growth conditions after 30’ and 60’ of induction of Cre (FX226 and FX227 dataset, respectively) and Cre-based Hi-SC2 profile of the 90’ time point of Figure 1 (FX224 dataset). A schematic representation of the site-specific recombination cassette for the Cre-activity reporter and Cre-based SC2 reporter is drawn in the left part of the panel. The excision of the Cre-activity reporter cassette reached an average of 80% after only 30’. This is much more than ***f_SC_*_2_** at all genomic positions after 30’, 60’ and 90’ of Cre production (Figure 1), in agreement with the hypothesis that replacement of one of the two SC2 reporter copies by a single SSR site mainly resulted from intermolecular recombination events. With the exception of a few loci, excision of the long cassette was constant along the genome, further indicating that Cre had a similar access to most chromosomal regions and was active at most positions. (B) Excision frequencies of the FX227 dataset versus the ***f_SC_*_2_** of the FX224 dataset. The horizontal and vertical dotted lines indicate the position of the median frequencies of excision frequencies in the FX227 data set and ***f_SC_*_2_** in the FX224 dataset, respectively. Long-distance variations in the frequency of excision of the SC2-reporter cassettes do not correlate with Cre-based ***f_SC_*_2_** variations.

**Figure S5.**
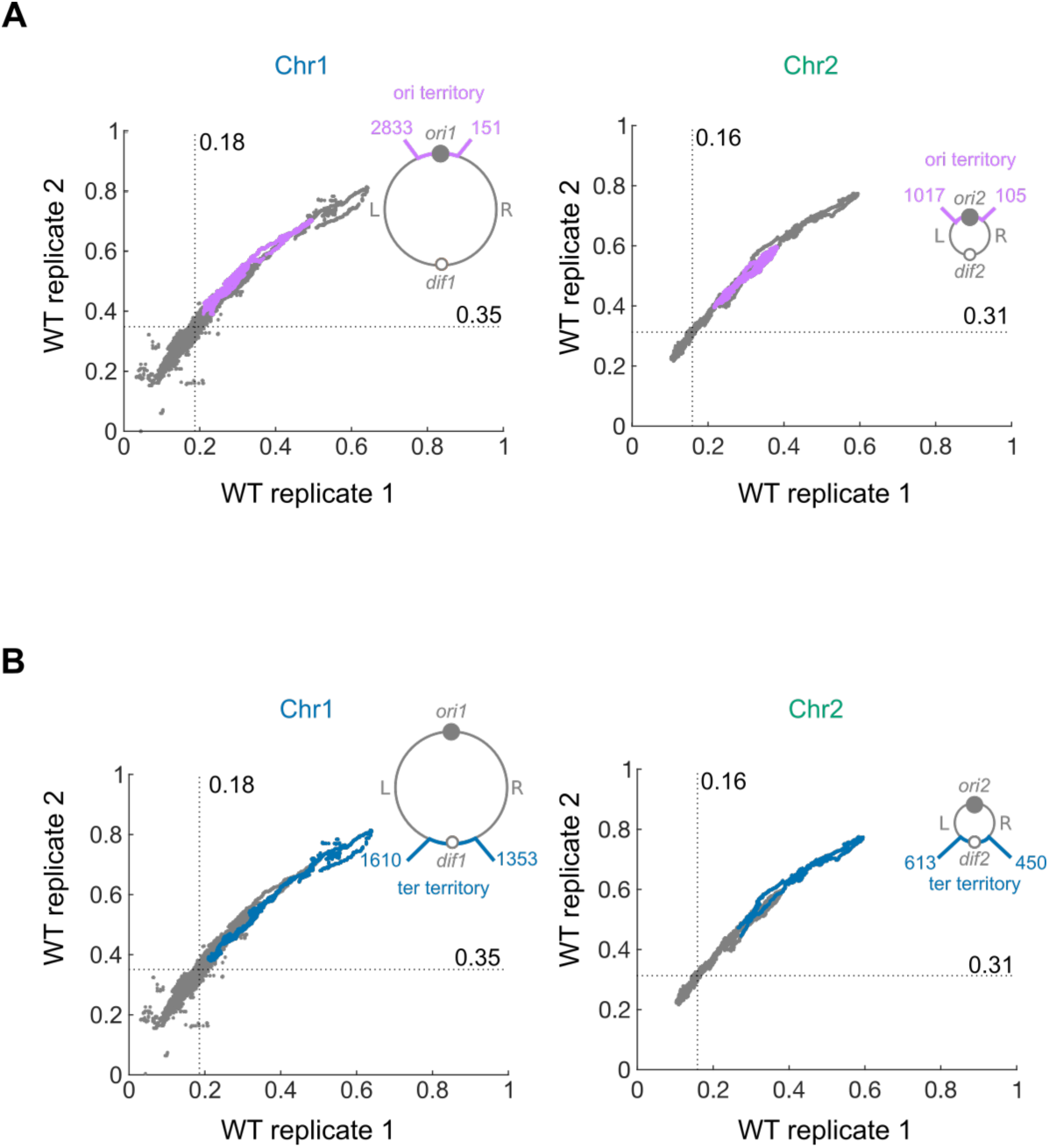
Large territories centred around the origin and terminus of replication of the two *V. cholerae* chromosomes are highly-cohesive. Cre-based ***f_SC_*_2_** of the FX239 dataset vs Cre-based ***f_SC_*_2_** of the FX224 dataset. The horizontal and vertical dotted lines indicate the position of the median ***f_SC_*_2_** in the FX239 and FX224 dataset, respectively. Positions belonging to the *ori1* and *ori2* territories (A) and *dif1* and *dif2* domains territories (B) are highlighted in pink and blue, respectively.

**Figure S6.**
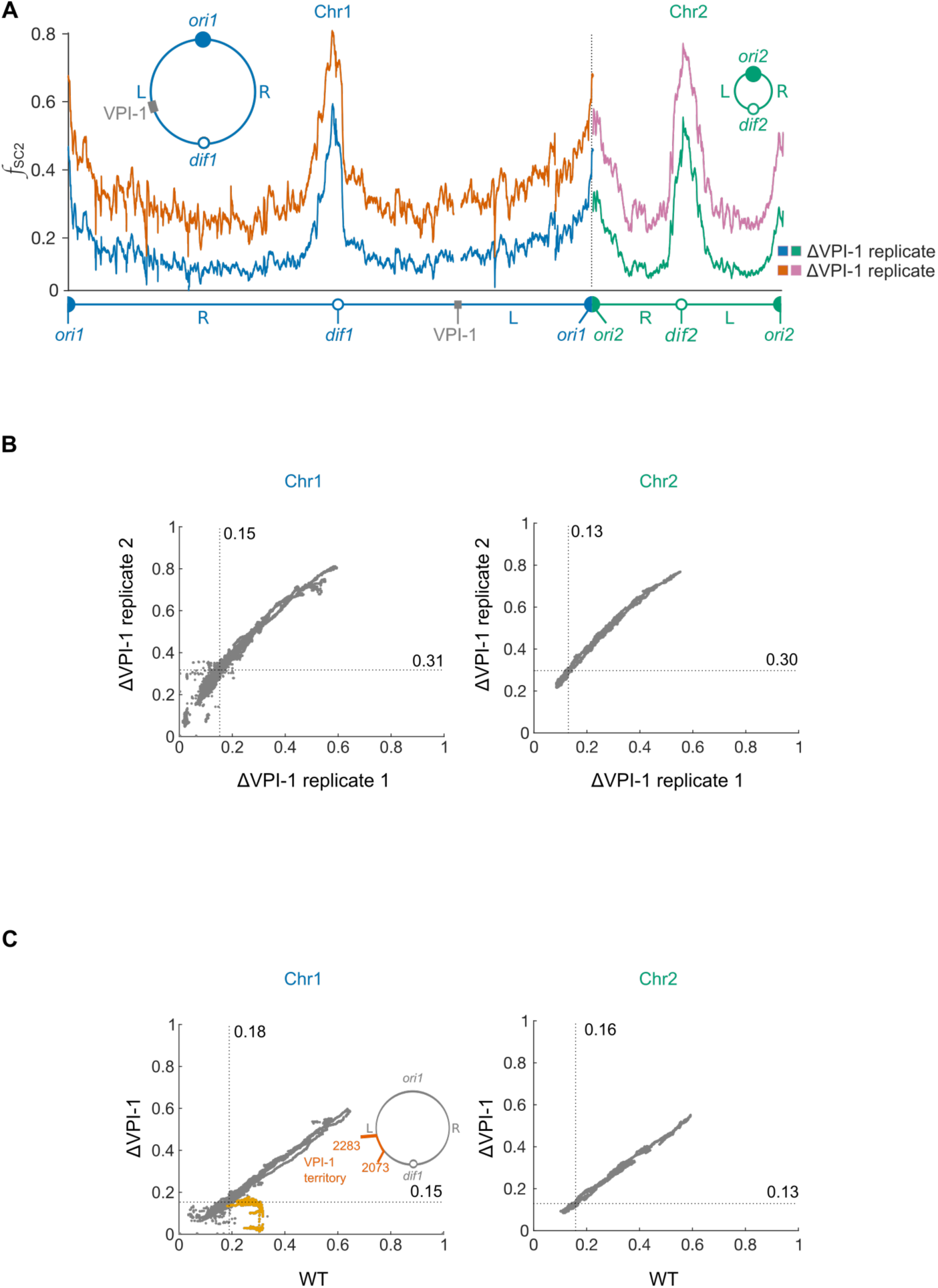
VPI-1 is responsible for the high cohesiveness of the VPI-1 domain. (A) Reproducibility of the Hi-SC2 profiles in ΔVPI-1 cells. Comparison of the 90’ time point of the Cre-based Hi-SC2 profile of Figure 2A (FX230 dataset) and the results obtained with an independent ΔVPI-1 transposition library at the same time point (FX243 dataset). (B) Cre-based ***f_SC_*_2_** from the FX230 dataset vs Cre-based ***f_SC_*_2_** from the FX243 dataset. The dotted horizontal and vertical lines indicate the position of the median ***f_SC_*_2_** in the FX243 and FX230 datasets, respectively. (C) Cre-based ***f_SC_*_2_** in ΔVPI-1 cells (FX230 dataset) vs Cre-based ***f_SC_*_2_** in wild-type cells (FX224 dataset). The dotted horizontal and vertical lines indicate the position of the median ***f_SC_*_2_** in the FX230 and FX224 datasets, respectively. The VPI-1 territory is indicated in orange. Loss of SC2 in the VPI-1 territory when VPI-1 is deleted is highlighted in orange (from position 2176653 to position 2217966).

**Figure S7.**
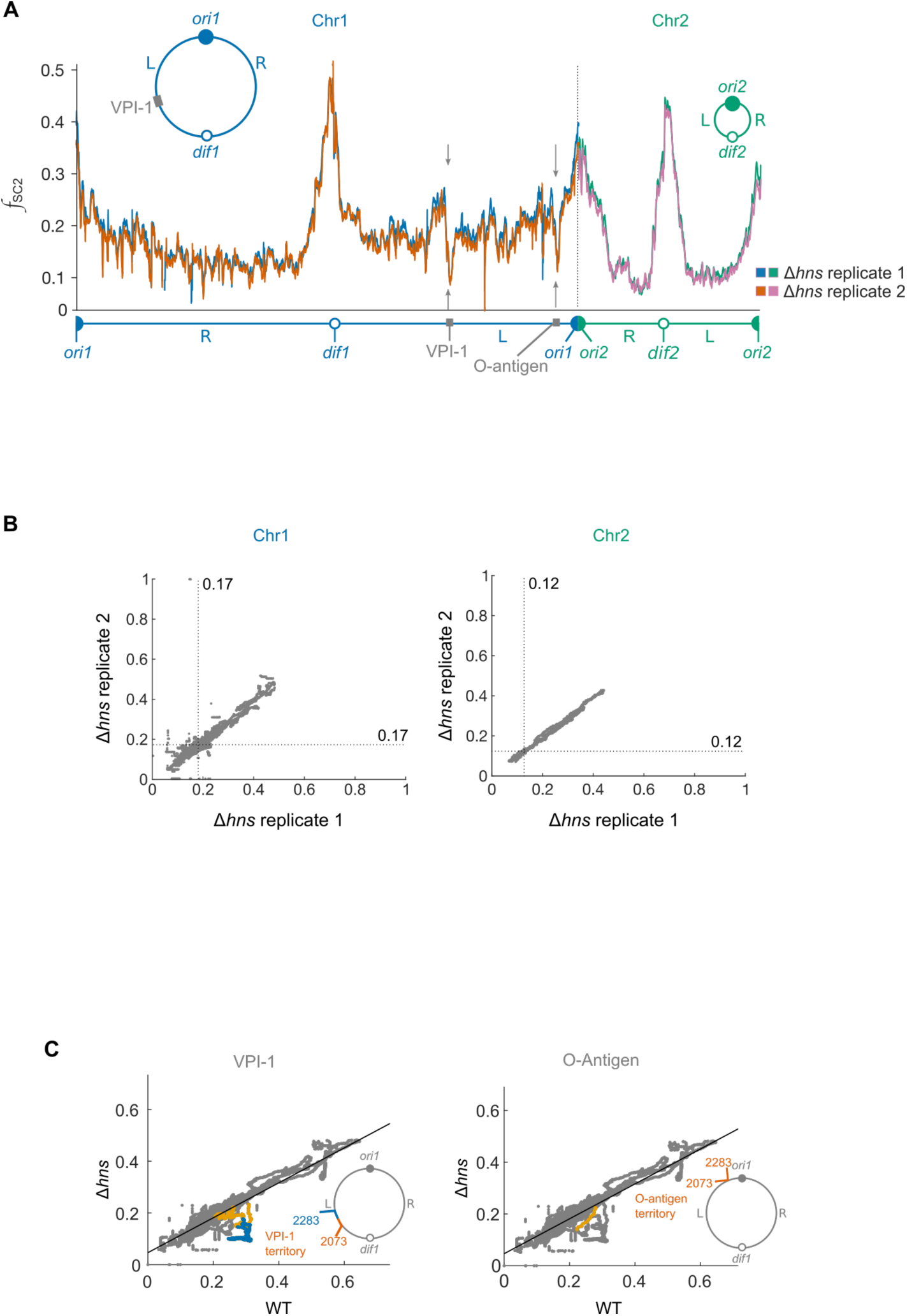
H-NS increases the cohesiveness of two regions implicated in *V. cholerae* virulence. (A) Reproducibility of the Hi-SC2 profiles in Δ*hns* strains. Comparison of the 90’ time point of the Cre-based Hi-SC2 profile of Figure 3A (FX215 dataset) and the results obtained with an independent cell library at the same time point (FX241 dataset). Grey arrows indicate regions of significantly lower SC2 frequencies in Δ*hns* cells. (B) Cre-based ***f_SC_*_2_** in the FX215 dataset vs Cre-based ***f_SC_*_2_** in the FX241 dataset. The dotted horizontal and vertical lines indicate the position of the median ***f_SC_*_2_** in the FX241 and FX215 datasets, respectively. (C) Cre-based ***f_SC_*_2_** in Δ*hns* cells (FX215 dataset) vs Cre-based ***f_SC_*_2_** in wild-type cells (FX224 dataset). Black line: linear regression of the Δ*hns* and WT data. Left panel: data points within VPI-1 and the 2160 to 2283 kbp origin-proximal part of the VPI-1 territory are highlighted in blue and orange, respectively; Right panel: data points within the 2825477 to 2839408 bp O-antigen region are highlighted in orange.

**Table S1.** List of Hi-SC2 experiments.

| Strain | Reporter | Clone number | Media | Cre induction time (min) | Dataset | Read number | Valid number of reads | Mapped insertions |
| --- | --- | --- | --- | --- | --- | --- | --- | --- |
| EEV29 | Xer-based SC2 reporter | 347,000 | M9 | NI | FX202 | 18,018,685 | 17,007,277 | 14,997,563 |
|  |  |  |  | 300 | FX205 | 17,066,405 | 16,494,988 | 14,637,494 |
|  | Cre-based SC2 reporter | 467,000 | M9 | NI | FX221 | 23,837,826 | 23,087,018 | 15,483,425 |
|  |  |  |  | 30 | FX222 | 27,374,911 | 26,589,027 | 19,411,784 |
|  |  |  |  | 60 | FX223 | 12,557,780 | 11,714,405 | 8,951,733 |
|  |  |  |  | 90 | FX224 | 26,430,938 | 25,864,693 | 20,245,240 |
|  | Cre-based SC2 reporter | 400,000 | M9 | NI | FX238 | 12,376,913 | 11,646,264 | 10,058,544 |
|  |  |  |  | 90 | FX239 | 15,941,629 | 15,500,388 | 13,497,374 |
|  | Cre-activity reporter | 569,000 | M9 | NI | FX225 | 14,627,990 | 14,308,341 | 10,540,592 |
|  |  |  |  | 30 | FX226 | 28,270,676 | 27,679,114 | 22,665,278 |
|  |  |  |  | 60 | FX227 | 26,683,039 | 25,608,567 | 21,300,506 |
|  | Cre-based SC2 reporter | 386,000 | LB | NI | FX126 | 11,593,777 | 11,393,652 | 10,287,107 |
|  |  |  |  | 60 | FX129 | 11,420,532 | 11,127,758 | 9,990,084 |
| EEV111 | Cre-based SC2 reporter | 431,000 | M9 | NI | FX229 | 26,947,308 | 26220063 | 10,713,596 |
|  |  |  |  | 90 | FX230 | 42,261,001 | 41,160,071 | 22,040,745 |
| EEV112 | Cre-based SC2 reporter | 476,000 | M9 | NI | FX242 | 14,106,539 | 13,763,589 | 11,036,874 |
|  |  |  |  | 90 | FX243 | 20,020,203 | 19,444,546 | 16,501,217 |
| EEV113 | Cre-based SC2 reporter | 607,000 | M9 | NI | FX214 | 31,709,306 | 30,847,276 | 27,607,681 |
|  |  |  |  | 90 | FX215 | 34,713,327 | 33,885,412 | 30,922,757 |
| EEV114 | Cre-based SC2 reporter | 431,000 | M9 | NI | FX240 | 20,439,647 | 19,598,167 | 18,179,412 |
|  |  |  |  | 90 | FX241 | 16,383,337 | 15,965,291 | 14,795,788 |

NI: Not induced. The Hi-SC2 Sequence data was deposited in the ArrayExpress Archive of Functional Genomics Data (https://www.ebi.ac.uk/arrayexpress/).

**Table S2.** Strains used in the study.

| Strain | Genotype/Description | Reference |
| --- | --- | --- |
| EEV29 | N16961 *ChapR ΔlacZ* P<sub>BAD</sub>-Cre-invP<sub>lac</sub>::FRT-*ble*-FRT, Zeo<sup>R</sup>, Gm<sup>R</sup> | This study |
| EEV111 | EEV29 ΔVPI-1::*bla*, Amp<sup>R</sup>, Zeo<sup>R</sup>, Gm<sup>R</sup> | This study |
| EEV112 | EEV29 ΔVPI-1::*bla*, Amp<sup>R</sup>, Zeo<sup>R</sup>, Gm<sup>R</sup> | This study |
| EEV113 | EEV29 Δ*hns*::*bla*, Amp<sup>R</sup>, Zeo<sup>R</sup>, Gm<sup>R</sup> | This study |
| EEV114 | EEV29 Δ*hns*::*bla*, Amp<sup>R</sup>, Zeo<sup>R</sup>, Gm<sup>R</sup> | This study |
| EEV136 | N16961 *ChapR ΔlacZ* P<sub>lac</sub>-*lacI*-RFP-*parB*<sub>PMT1</sub>-YGFP + LacOarray-*aph* on chromosome 1 at position 2.043 Mb + tet'-*parS*<sub>PMT1</sub>-*cat*-tet inserted on chromosome 1 at position 2.12 MB, Kan<sup>R</sup>, Cm<sup>R</sup>, Gm<sup>R</sup> | This study |
| EEV139 | N16961 *ChapR ΔlacZ* P<sub>lac</sub>-*lacI*-RFP-*parB*<sub>PMT1</sub>-YGFP + LacO array-*aph* on chromosome 1 at position 2.043 Mb + tet'-*parS*<sub>PMT1</sub>-*cat*-tet inserted on chromosome 1 at position 2.12 Mb ΔVPI-1::*bla*, Kan<sup>R</sup>, Cm<sup>R</sup>, Gm<sup>R</sup>, Amp<sup>R</sup> | This study |
| EEV172 | N16961 *ChapR ΔlacZ* P<sub>lac</sub>-*lacI*-RFP-*parB*<sub>PMT1</sub>-YGFP + LacO array-*aph* on chromosome 1 at position 0.85 Mb + tet'-*parS*<sub>PMT1</sub>-*cat*-tet inserted on chromosome 1 at position 0.814 MB, Kan<sup>R</sup>, Cm<sup>R</sup>, Gm<sup>R</sup> | This study |
| GDV28 | N16961 *ChapR ΔlacZ* P<sub>BAD</sub>-*xerC*-invP<sub>lac</sub>-*aadA1* Spec<sup>R</sup>, Gm<sup>R</sup> | (Demarre et al., 2014) |
| DH5α | DH5α λpir | Lab stock |
| β2163 | (F<sup>-</sup>) RP4-2-Tc::Mu DAP<sup>-</sup> Kan<sup>R</sup> | (Demarre et al., 2005) |
| FCV14 | DH5α λpir Δ*xerC* | Lab stock |
| EE6 | β2163 Δ*xerC* Kan<sup>R</sup> Spec<sup>R</sup> | This study |
| EE13 | EE6/ pEE20 | This study |
| EE14 | β2163/ pEE22 | This study |
| EE15 | β2163/ pEE56 | This study |

Zeo: Zeocin; Gm: Gentamicin; Amp: Ampicilin; Kan: Kanamycin; Spec: Spectinomycin; Cm: Chloramphenicol

**Table S3.** Plasmids used in the study.

| Plasmid | Derived from | Description | Reference |
| --- | --- | --- | --- |
| pEE20 | pSW23T | Ori R6K Cm<sup>R</sup> *magellan5-Mmel* (Kan<sup>R</sup>). Rd2SP sequence. Xer-based SC2 reporter (27 bp between *dif1* sites). | This study |
| pEE22 | pSW23T | Ori R6K Cm<sup>R</sup> *magellan5-Mmel* (Kan<sup>R</sup>). Rd2SP sequence. Cre-based SC2 reporter (20 bp between *loxP* sites). | This study |
| pEE23 | pUC18 | Plasmid for natural transformation carrying P<sub>BAD</sub> –Cre-*invP<sub>lac</sub>::FRT-ble-FRT* between two homologous regions of *V. cholerae lacZ* , Amp<sup>R</sup>, Zeo<sup>R</sup> | This study |
| pEE44 | pUC18 | Plasmid for natural transformation carrying FRT-*bla*-FRT between homologous regions of *V. cholerae* upstream and downstream of VPI-1 Cm<sup>R</sup>, Amp<sup>R</sup> | This study |
| pEE46 | pUC18 | Plasmid for natural transformation carrying FRT-*bla*-FRT between homologous regions of *V. cholerae* upstream and downstream of *hns* Cm<sup>R</sup>, Amp<sup>R</sup> | This study |
| pEE56 | pSW23T | Ori R6K Cm<sup>R</sup> *magellan5-Mmel* (Kan<sup>R</sup>). Rd2SP sequence. Cre-activity reporter (205 bp between *loxP* sites). | This study |
| pSW23T | - | Ori R6K Cm<sup>R</sup> conjugative vector | (Demarre et al., 2014) |

**Table S4.**
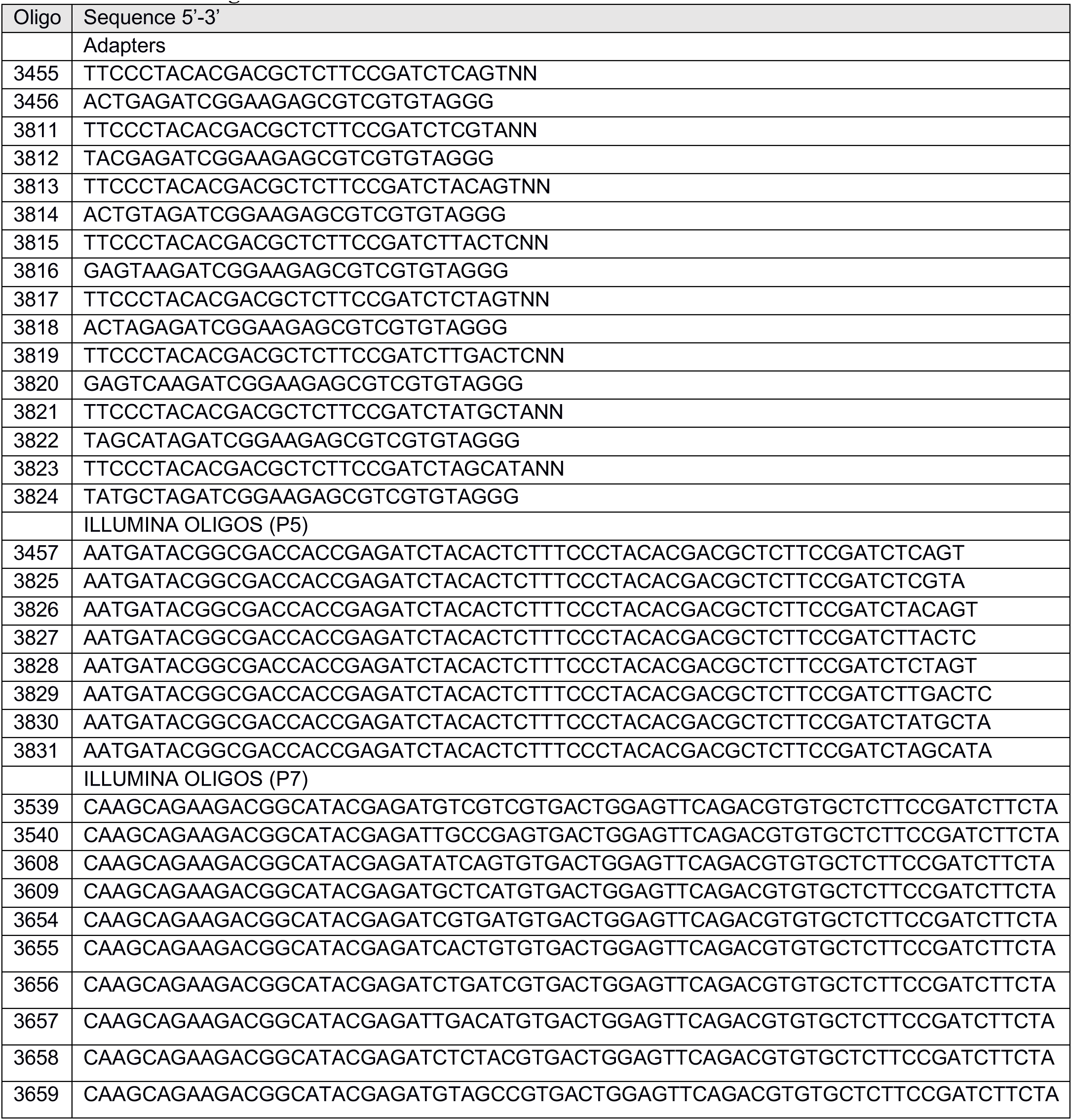
Oligonucleotides.

**Table S5.**
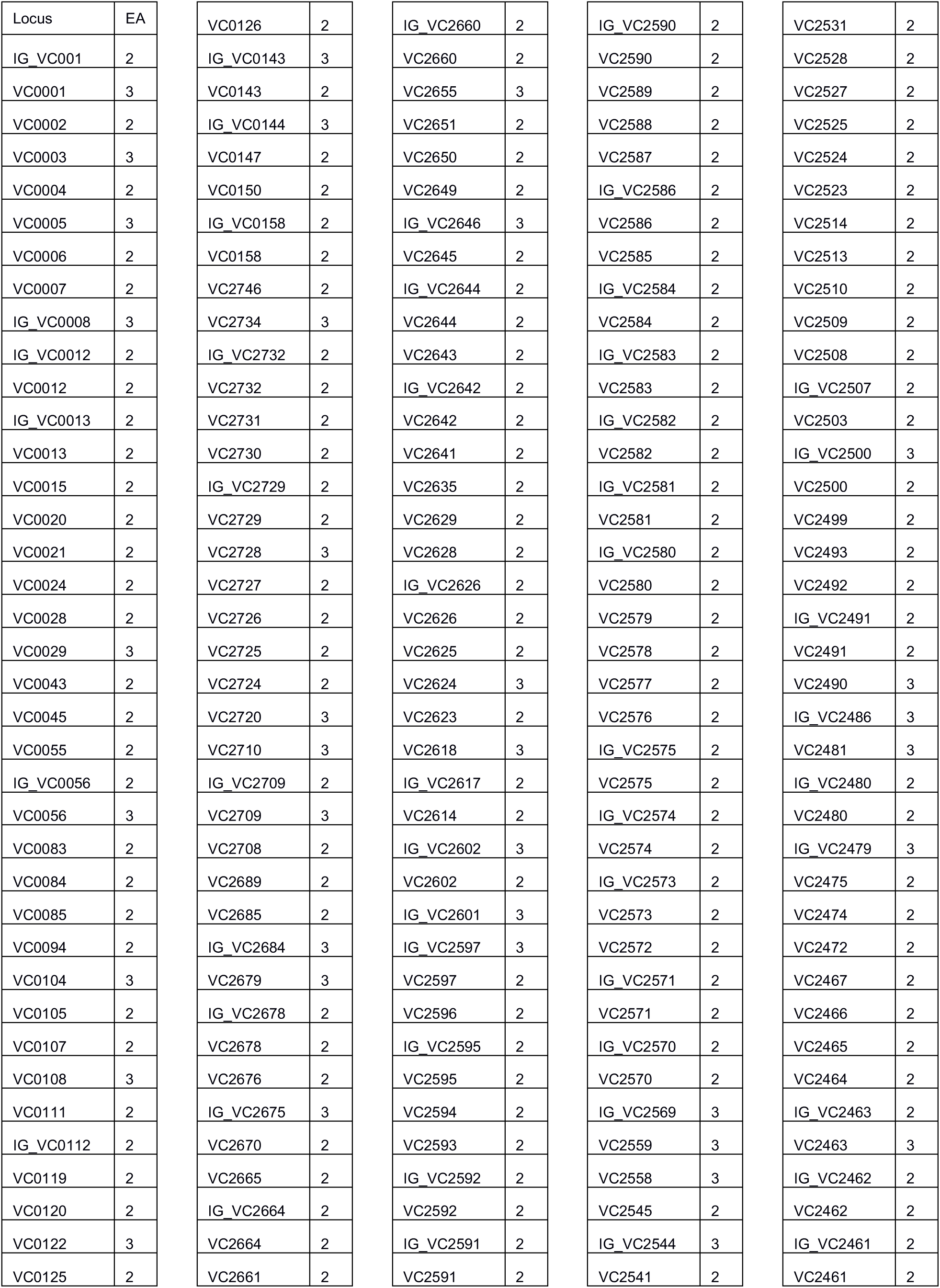
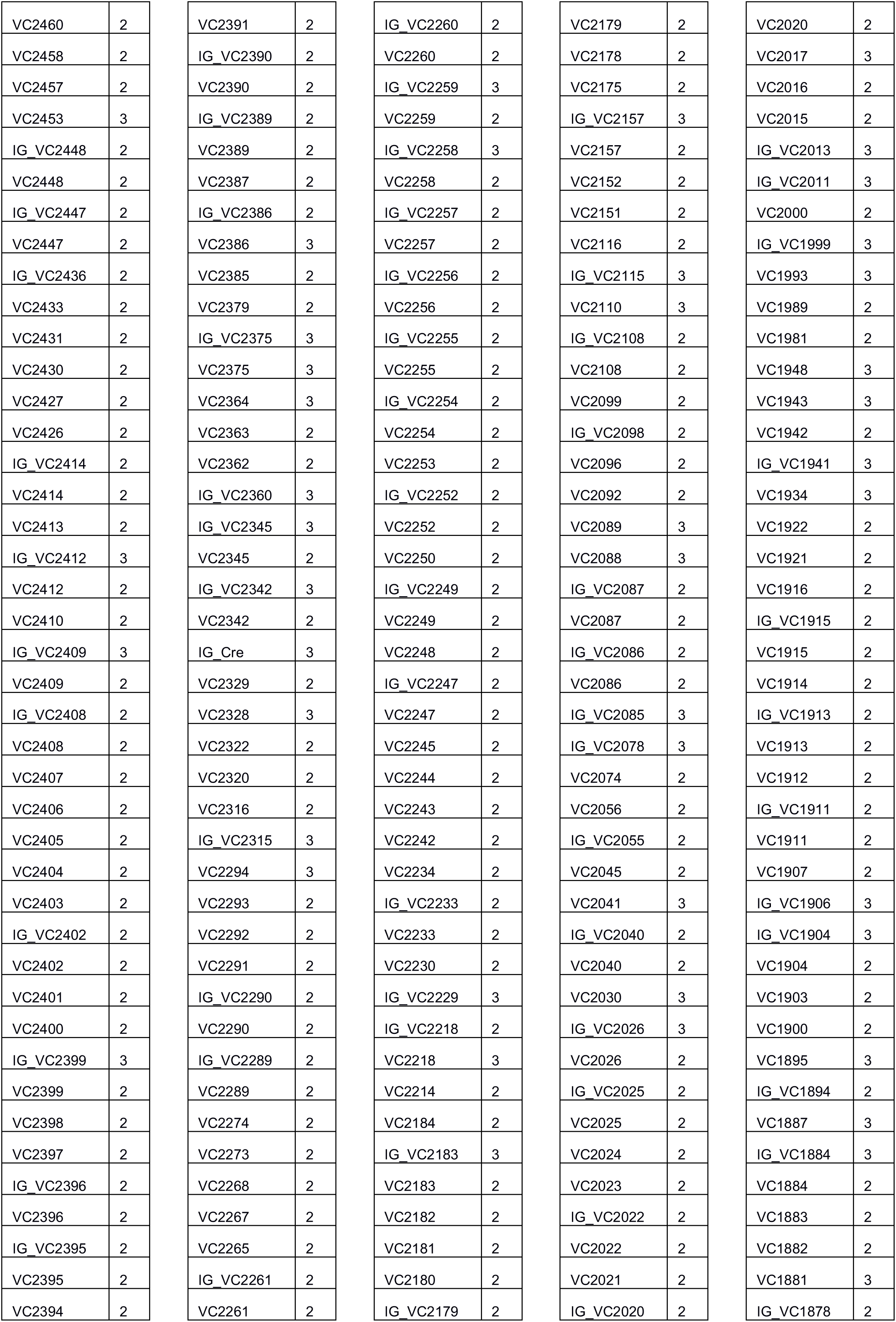
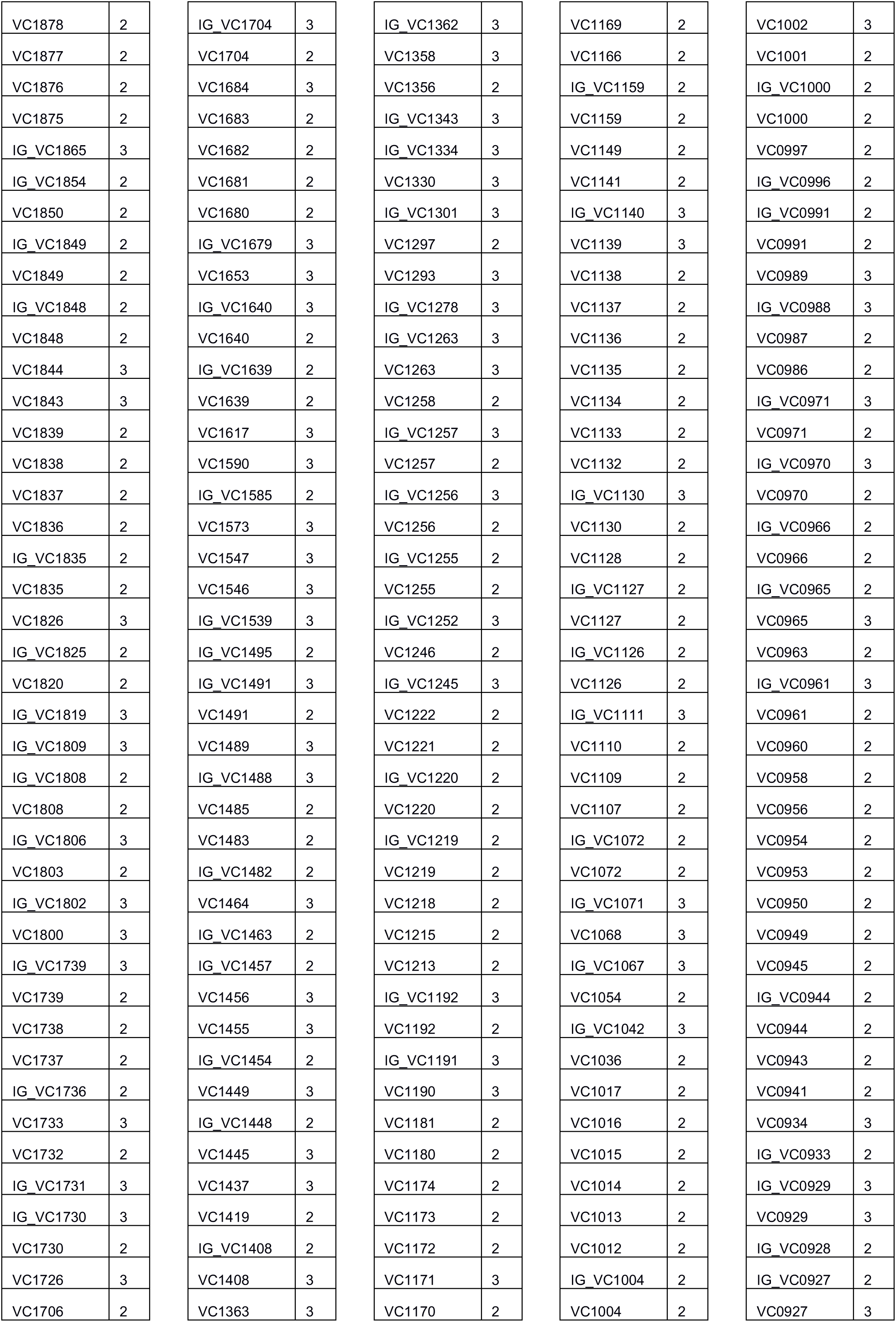
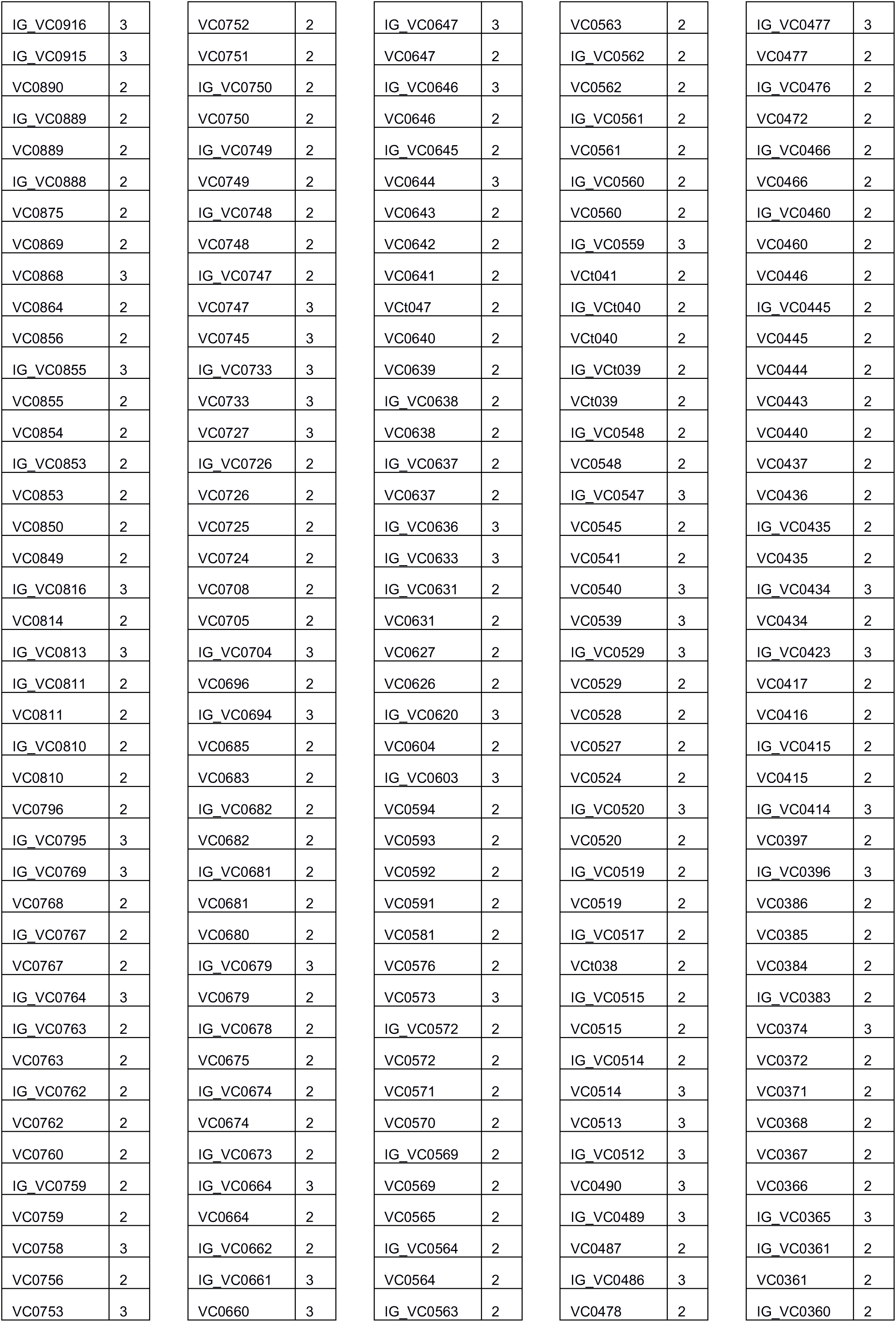
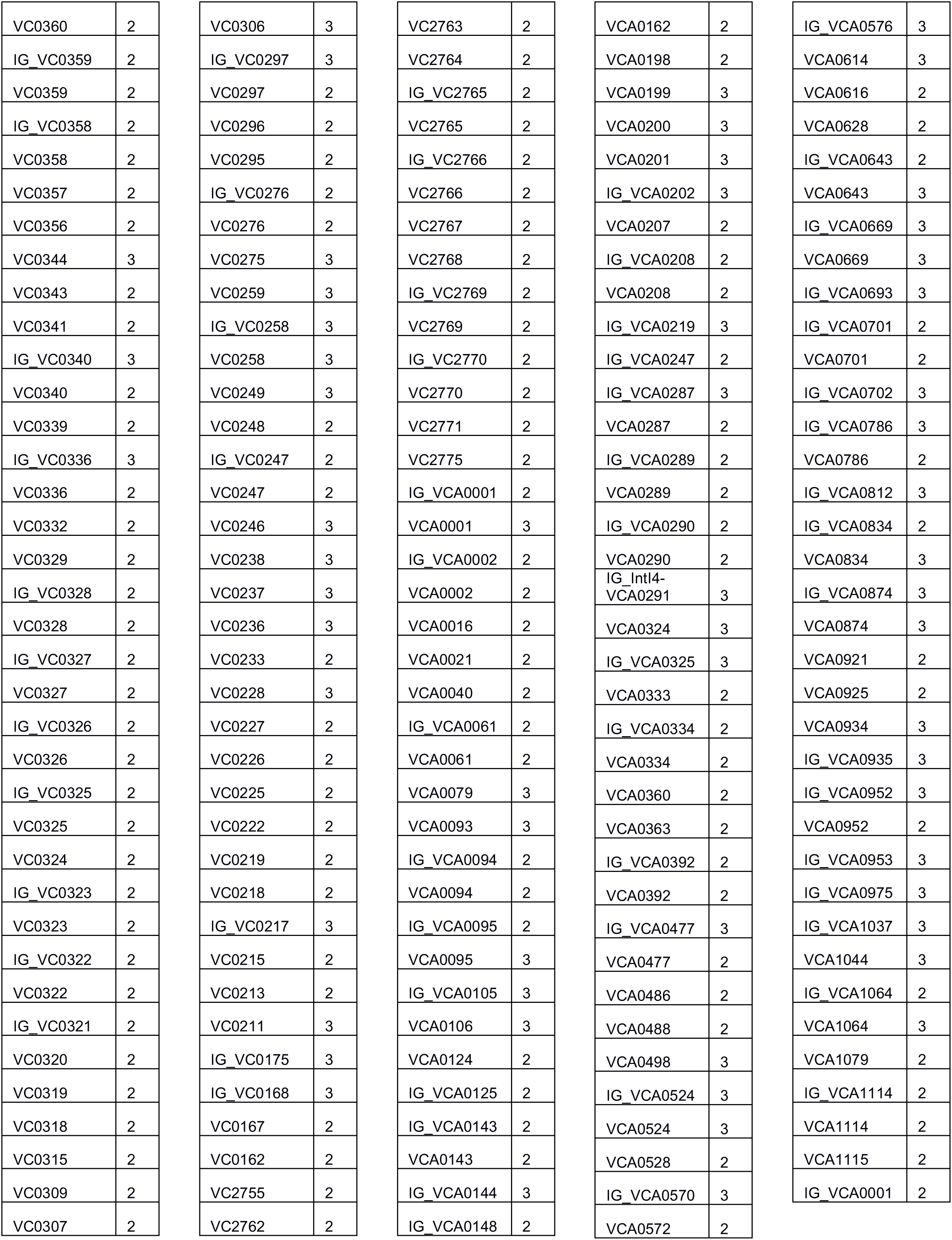
Essential genes and intergenic regions. EA: EL-ARTIST; 2: locus essential for optimal growth; 3: locus domain essential; IG: Intergenic region.

|  |  |  |  |  |  |  |  |  |  |
| --- | --- | --- | --- | --- | --- | --- | --- | --- | --- |
| Locus | EA | VC0126 | 2 | IG_VC2660 | 2 | IG_VC2590 | 2 | VC2531 | 2 |
| IG_VC001 | 2 | IG_VC0143 | 3 | VC2660 | 2 | VC2590 | 2 | VC2528 | 2 |
| VC0001 | 3 | VC0143 | 2 | VC2655 | 3 | VC2589 | 2 | VC2527 | 2 |
| VC0002 | 2 | IG_VC0144 | 3 | VC2651 | 2 | VC2588 | 2 | VC2525 | 2 |
| VC0003 | 3 | VC0147 | 2 | VC2650 | 2 | VC2587 | 2 | VC2524 | 2 |
| VC0004 | 2 | VC0150 | 2 | VC2649 | 2 | IG_VC2586 | 2 | VC2523 | 2 |
| VC0005 | 3 | IG_VC0158 | 2 | IG_VC2646 | 3 | VC2586 | 2 | VC2514 | 2 |
| VC0006 | 2 | VC0158 | 2 | VC2645 | 2 | VC2585 | 2 | VC2513 | 2 |
| VC0007 | 2 | VC2746 | 2 | IG_VC2644 | 2 | IG_VC2584 | 2 | VC2510 | 2 |
| IG_VC0008 | 3 | VC2734 | 3 | VC2644 | 2 | VC2584 | 2 | VC2509 | 2 |
| IG_VC0012 | 2 | IG_VC2732 | 2 | VC2643 | 2 | IG_VC2583 | 2 | VC2508 | 2 |
| VC0012 | 2 | VC2732 | 2 | IG_VC2642 | 2 | VC2583 | 2 | IG_VC2507 | 2 |
| IG_VC0013 | 2 | VC2731 | 2 | VC2642 | 2 | IG_VC2582 | 2 | VC2503 | 2 |
| VC0013 | 2 | VC2730 | 2 | VC2641 | 2 | VC2582 | 2 | IG_VC2500 | 3 |
| VC0015 | 2 | IG_VC2729 | 2 | VC2635 | 2 | IG_VC2581 | 2 | VC2500 | 2 |
| VC0020 | 2 | VC2729 | 2 | VC2629 | 2 | VC2581 | 2 | VC2499 | 2 |
| VC0021 | 2 | VC2728 | 3 | VC2628 | 2 | IG_VC2580 | 2 | VC2493 | 2 |
| VC0024 | 2 | VC2727 | 2 | IG_VC2626 | 2 | VC2580 | 2 | VC2492 | 2 |
| VC0028 | 2 | VC2726 | 2 | VC2626 | 2 | VC2579 | 2 | IG_VC2491 | 2 |
| VC0029 | 3 | VC2725 | 2 | VC2625 | 2 | VC2578 | 2 | VC2491 | 2 |
| VC0043 | 2 | VC2724 | 2 | VC2624 | 3 | VC2577 | 2 | VC2490 | 3 |
| VC0045 | 2 | VC2720 | 3 | VC2623 | 2 | VC2576 | 2 | IG_VC2486 | 3 |
| VC0055 | 2 | VC2710 | 3 | VC2618 | 3 | IG_VC2575 | 2 | VC2481 | 3 |
| IG_VC0056 | 2 | IG_VC2709 | 2 | IG_VC2617 | 2 | VC2575 | 2 | IG_VC2480 | 2 |
| VC0056 | 3 | VC2709 | 3 | VC2614 | 2 | IG_VC2574 | 2 | VC2480 | 2 |
| VC0083 | 2 | VC2708 | 2 | IG_VC2602 | 3 | VC2574 | 2 | IG_VC2479 | 3 |
| VC0084 | 2 | VC2689 | 2 | VC2602 | 2 | IG_VC2573 | 2 | VC2475 | 2 |
| VC0085 | 2 | VC2685 | 2 | IG_VC2601 | 3 | VC2573 | 2 | VC2474 | 2 |
| VC0094 | 2 | IG_VC2684 | 3 | IG_VC2597 | 3 | VC2572 | 2 | VC2472 | 2 |
| VC0104 | 3 | VC2679 | 3 | VC2597 | 2 | IG_VC2571 | 2 | VC2467 | 2 |
| VC0105 | 2 | IG_VC2678 | 2 | VC2596 | 2 | VC2571 | 2 | VC2466 | 2 |
| VC0107 | 2 | VC2678 | 2 | IG_VC2595 | 2 | IG_VC2570 | 2 | VC2465 | 2 |
| VC0108 | 3 | VC2676 | 2 | VC2595 | 2 | VC2570 | 2 | VC2464 | 2 |
| VC0111 | 2 | IG_VC2675 | 3 | VC2594 | 2 | IG_VC2569 | 3 | IG_VC2463 | 2 |
| IG_VC0112 | 2 | VC2670 | 2 | VC2593 | 2 | VC2559 | 3 | VC2463 | 3 |
| VC0119 | 2 | VC2665 | 2 | IG_VC2592 | 2 | VC2558 | 3 | IG_VC2462 | 2 |
| VC0120 | 2 | IG_VC2664 | 2 | VC2592 | 2 | VC2545 | 2 | VC2462 | 2 |
| VC0122 | 3 | VC2664 | 2 | IG_VC2591 | 2 | IG_VC2544 | 3 | IG_VC2461 | 2 |
| VC0125 | 2 | VC2661 | 2 | VC2591 | 2 | VC2541 | 2 | VC2461 | 2 |

|  |  |
| --- | --- |
| VC2460 | 2 |
| VC2458 | 2 |
| VC2457 | 2 |
| VC2453 | 3 |
| IG_VC2448 | 2 |
| VC2448 | 2 |
| IG_VC2447 | 2 |
| VC2447 | 2 |
| IG_VC2436 | 2 |
| VC2433 | 2 |
| VC2431 | 2 |
| VC2430 | 2 |
| VC2427 | 2 |
| VC2426 | 2 |
| IG_VC2414 | 2 |
| VC2414 | 2 |
| VC2413 | 2 |
| IG_VC2412 | 3 |
| VC2412 | 2 |
| VC2410 | 2 |
| IG_VC2409 | 3 |
| VC2409 | 2 |
| IG_VC2408 | 2 |
| VC2408 | 2 |
| VC2407 | 2 |
| VC2406 | 2 |
| VC2405 | 2 |
| VC2404 | 2 |
| VC2403 | 2 |
| IG_VC2402 | 2 |
| VC2402 | 2 |
| VC2401 | 2 |
| VC2400 | 2 |
| IG_VC2399 | 3 |
| VC2399 | 2 |
| VC2398 | 2 |
| VC2397 | 2 |
| IG_VC2396 | 2 |
| VC2396 | 2 |
| IG_VC2395 | 2 |
| VC2395 | 2 |
| VC2394 | 2 |
| VC2391 | 2 |
| IG_VC2390 | 2 |
| VC2390 | 2 |
| IG_VC2389 | 2 |
| VC2389 | 2 |
| VC2387 | 2 |
| IG_VC2386 | 2 |
| VC2386 | 3 |
| VC2385 | 2 |
| VC2379 | 2 |
| IG_VC2375 | 3 |
| VC2375 | 3 |
| VC2364 | 3 |
| VC2363 | 2 |
| VC2362 | 2 |
| IG_VC2360 | 3 |
| IG_VC2345 | 3 |
| VC2345 | 2 |
| IG_VC2342 | 3 |
| VC2342 | 2 |
| IG_Cre | 3 |
| VC2329 | 2 |
| VC2328 | 3 |
| VC2322 | 2 |
| VC2320 | 2 |
| VC2316 | 2 |
| IG_VC2315 | 3 |
| VC2294 | 3 |
| VC2293 | 2 |
| VC2292 | 2 |
| VC2291 | 2 |
| IG_VC2290 | 2 |
| VC2290 | 2 |
| IG_VC2289 | 2 |
| VC2289 | 2 |
| VC2274 | 2 |
| VC2273 | 2 |
| VC2268 | 2 |
| VC2267 | 2 |
| VC2265 | 2 |
| IG_VC2261 | 2 |
| VC2261 | 2 |
| IG_VC2260 | 2 |
| VC2260 | 2 |
| IG_VC2259 | 3 |
| VC2259 | 2 |
| IG_VC2258 | 3 |
| VC2258 | 2 |
| IG_VC2257 | 2 |
| VC2257 | 2 |
| IG_VC2256 | 2 |
| VC2256 | 2 |
| IG_VC2255 | 2 |
| VC2255 | 2 |
| IG_VC2254 | 2 |
| VC2254 | 2 |
| VC2253 | 2 |
| IG_VC2252 | 2 |
| VC2252 | 2 |
| VC2250 | 2 |
| IG_VC2249 | 2 |
| VC2249 | 2 |
| VC2248 | 2 |
| IG_VC2247 | 2 |
| VC2247 | 2 |
| VC2245 | 2 |
| VC2244 | 2 |
| VC2243 | 2 |
| VC2242 | 2 |
| VC2234 | 2 |
| IG_VC2233 | 2 |
| VC2233 | 2 |
| VC2230 | 2 |
| IG_VC2229 | 3 |
| IG_VC2218 | 2 |
| VC2218 | 3 |
| VC2214 | 2 |
| VC2184 | 2 |
| IG_VC2183 | 3 |
| VC2183 | 2 |
| VC2182 | 2 |
| VC2181 | 2 |
| VC2180 | 2 |
| IG_VC2179 | 2 |
| VC2179 | 2 |
| VC2178 | 2 |
| VC2175 | 2 |
| IG_VC2157 | 3 |
| VC2157 | 2 |
| VC2152 | 2 |
| VC2151 | 2 |
| VC2116 | 2 |
| IG_VC2115 | 3 |
| VC2110 | 3 |
| IG_VC2108 | 2 |
| VC2108 | 2 |
| VC2099 | 2 |
| IG_VC2098 | 2 |
| VC2096 | 2 |
| VC2092 | 2 |
| VC2089 | 3 |
| VC2088 | 3 |
| IG_VC2087 | 2 |
| VC2087 | 2 |
| IG_VC2086 | 2 |
| VC2086 | 2 |
| IG_VC2085 | 3 |
| IG_VC2078 | 3 |
| VC2074 | 2 |
| VC2056 | 2 |
| IG_VC2055 | 2 |
| VC2045 | 2 |
| VC2041 | 3 |
| IG_VC2040 | 2 |
| VC2040 | 2 |
| VC2030 | 3 |
| IG_VC2026 | 3 |
| VC2026 | 2 |
| IG_VC2025 | 2 |
| VC2025 | 2 |
| VC2024 | 2 |
| VC2023 | 2 |
| IG_VC2022 | 2 |
| VC2022 | 2 |
| VC2021 | 2 |
| IG_VC2020 | 2 |
| VC2020 | 2 |
| VC2017 | 3 |
| VC2016 | 2 |
| VC2015 | 2 |
| IG_VC2013 | 3 |
| IG_VC2011 | 3 |
| VC2000 | 2 |
| IG_VC1999 | 3 |
| VC1993 | 3 |
| VC1989 | 2 |
| VC1981 | 2 |
| VC1948 | 3 |
| VC1943 | 3 |
| VC1942 | 2 |
| IG_VC1941 | 3 |
| VC1934 | 3 |
| VC1922 | 2 |
| VC1921 | 2 |
| VC1916 | 2 |
| IG_VC1915 | 2 |
| VC1915 | 2 |
| VC1914 | 2 |
| IG_VC1913 | 2 |
| VC1913 | 2 |
| VC1912 | 2 |
| IG_VC1911 | 2 |
| VC1911 | 2 |
| VC1907 | 2 |
| IG_VC1906 | 3 |
| IG_VC1904 | 3 |
| VC1904 | 2 |
| VC1903 | 2 |
| VC1900 | 2 |
| VC1895 | 3 |
| IG_VC1894 | 2 |
| VC1887 | 3 |
| IG_VC1884 | 3 |
| VC1884 | 2 |
| VC1883 | 2 |
| VC1882 | 2 |
| VC1881 | 3 |
| IG_VC1878 | 2 |
| VC1878 | 2 |
| VC1877 | 2 |
| VC1876 | 2 |
| VC1875 | 2 |
| IG_VC1865 | 3 |
| IG_VC1854 | 2 |
| VC1850 | 2 |
| IG_VC1849 | 2 |
| VC1849 | 2 |
| IG_VC1848 | 2 |
| VC1848 | 2 |
| VC1844 | 3 |
| VC1843 | 3 |
| VC1839 | 2 |
| VC1838 | 2 |
| VC1837 | 2 |
| VC1836 | 2 |
| IG_VC1835 | 2 |
| VC1835 | 2 |
| VC1826 | 3 |
| IG_VC1825 | 2 |
| VC1820 | 2 |
| IG_VC1819 | 3 |
| IG_VC1809 | 3 |
| IG_VC1808 | 2 |
| VC1808 | 2 |
| IG_VC1806 | 3 |
| VC1803 | 2 |
| IG_VC1802 | 3 |
| VC1800 | 3 |
| IG_VC1739 | 3 |
| VC1739 | 2 |
| VC1738 | 2 |
| VC1737 | 2 |
| IG_VC1736 | 2 |
| VC1733 | 3 |
| VC1732 | 2 |
| IG_VC1731 | 3 |
| IG_VC1730 | 3 |
| VC1730 | 2 |
| VC1726 | 3 |
| VC1706 | 2 |
| IG_VC1704 | 3 |
| VC1704 | 2 |
| VC1684 | 3 |
| VC1683 | 2 |
| VC1682 | 2 |
| VC1681 | 2 |
| VC1680 | 2 |
| IG_VC1679 | 3 |
| VC1653 | 3 |
| IG_VC1640 | 3 |
| VC1640 | 2 |
| IG_VC1639 | 2 |
| VC1639 | 2 |
| VC1617 | 3 |
| VC1590 | 3 |
| IG_VC1585 | 2 |
| VC1573 | 3 |
| VC1547 | 3 |
| VC1546 | 3 |
| IG_VC1539 | 3 |
| IG_VC1495 | 2 |
| IG_VC1491 | 3 |
| VC1491 | 2 |
| VC1489 | 3 |
| IG_VC1488 | 3 |
| VC1485 | 2 |
| VC1483 | 2 |
| IG_VC1482 | 2 |
| VC1464 | 3 |
| IG_VC1463 | 2 |
| IG_VC1457 | 2 |
| VC1456 | 3 |
| VC1455 | 3 |
| IG_VC1454 | 2 |
| VC1449 | 3 |
| IG_VC1448 | 2 |
| VC1445 | 3 |
| VC1437 | 3 |
| VC1419 | 2 |
| IG_VC1408 | 2 |
| VC1408 | 3 |
| VC1363 | 3 |
| IG_VC1362 | 3 |
| VC1358 | 3 |
| VC1356 | 2 |
| IG_VC1343 | 3 |
| IG_VC1334 | 3 |
| VC1330 | 3 |
| IG_VC1301 | 3 |
| VC1297 | 2 |
| VC1293 | 3 |
| IG_VC1278 | 3 |
| IG_VC1263 | 3 |
| VC1263 | 3 |
| VC1258 | 2 |
| IG_VC1257 | 3 |
| VC1257 | 2 |
| IG_VC1256 | 3 |
| VC1256 | 2 |
| IG_VC1255 | 2 |
| VC1255 | 2 |
| IG_VC1252 | 3 |
| VC1246 | 2 |
| IG_VC1245 | 3 |
| VC1222 | 2 |
| VC1221 | 2 |
| IG_VC1220 | 2 |
| VC1220 | 2 |
| IG_VC1219 | 2 |
| VC1219 | 2 |
| VC1218 | 2 |
| VC1215 | 2 |
| VC1213 | 2 |
| IG_VC1192 | 3 |
| VC1192 | 2 |
| IG_VC1191 | 3 |
| VC1190 | 3 |
| VC1181 | 2 |
| VC1180 | 2 |
| VC1174 | 2 |
| VC1173 | 2 |
| VC1172 | 2 |
| VC1171 | 3 |
| VC1170 | 2 |
| VC1169 | 2 |
| VC1166 | 2 |
| IG_VC1159 | 2 |
| VC1159 | 2 |
| VC1149 | 2 |
| VC1141 | 2 |
| IG_VC1140 | 3 |
| VC1139 | 3 |
| VC1138 | 2 |
| VC1137 | 2 |
| VC1136 | 2 |
| VC1135 | 2 |
| VC1134 | 2 |
| VC1133 | 2 |
| VC1132 | 2 |
| IG_VC1130 | 3 |
| VC1130 | 2 |
| VC1128 | 2 |
| IG_VC1127 | 2 |
| VC1127 | 2 |
| IG_VC1126 | 2 |
| VC1126 | 2 |
| IG_VC1111 | 3 |
| VC1110 | 2 |
| VC1109 | 2 |
| VC1107 | 2 |
| IG_VC1072 | 2 |
| VC1072 | 2 |
| IG_VC1071 | 3 |
| VC1068 | 3 |
| IG_VC1067 | 3 |
| VC1054 | 2 |
| IG_VC1042 | 3 |
| VC1036 | 2 |
| VC1017 | 2 |
| VC1016 | 2 |
| VC1015 | 2 |
| VC1014 | 2 |
| VC1013 | 2 |
| VC1012 | 2 |
| IG_VC1004 | 2 |
| VC1004 | 2 |
| VC1002 | 3 |
| VC1001 | 2 |
| IG_VC1000 | 2 |
| VC1000 | 2 |
| VC0997 | 2 |
| IG_VC0996 | 2 |
| IG_VC0991 | 2 |
| VC0991 | 2 |
| VC0989 | 3 |
| IG_VC0988 | 3 |
| VC0987 | 2 |
| VC0986 | 2 |
| IG_VC0971 | 3 |
| VC0971 | 2 |
| IG_VC0970 | 3 |
| VC0970 | 2 |
| IG_VC0966 | 2 |
| VC0966 | 2 |
| IG_VC0965 | 2 |
| VC0965 | 3 |
| VC0963 | 2 |
| IG_VC0961 | 3 |
| VC0961 | 2 |
| VC0960 | 2 |
| VC0958 | 2 |
| VC0956 | 2 |
| VC0954 | 2 |
| VC0953 | 2 |
| VC0950 | 2 |
| VC0949 | 2 |
| VC0945 | 2 |
| IG_VC0944 | 2 |
| VC0944 | 2 |
| VC0943 | 2 |
| VC0941 | 2 |
| VC0934 | 3 |
| IG_VC0933 | 2 |
| IG_VC0929 | 3 |
| VC0929 | 3 |
| IG_VC0928 | 2 |
| IG_VC0927 | 2 |
| VC0927 | 3 |
| IG_VC0916 | 3 |
| IG_VC0915 | 3 |
| VC0890 | 2 |
| IG_VC0889 | 2 |
| VC0889 | 2 |
| IG_VC0888 | 2 |
| VC0875 | 2 |
| VC0869 | 2 |
| VC0868 | 3 |
| VC0864 | 2 |
| VC0856 | 2 |
| IG_VC0855 | 3 |
| VC0855 | 2 |
| VC0854 | 2 |
| IG_VC0853 | 2 |
| VC0853 | 2 |
| VC0850 | 2 |
| VC0849 | 2 |
| IG_VC0816 | 3 |
| VC0814 | 2 |
| IG_VC0813 | 3 |
| IG_VC0811 | 2 |
| VC0811 | 2 |
| IG_VC0810 | 2 |
| VC0810 | 2 |
| VC0796 | 2 |
| IG_VC0795 | 3 |
| IG_VC0769 | 3 |
| VC0768 | 2 |
| IG_VC0767 | 2 |
| VC0767 | 2 |
| IG_VC0764 | 3 |
| IG_VC0763 | 2 |
| VC0763 | 2 |
| IG_VC0762 | 2 |
| VC0762 | 2 |
| VC0760 | 2 |
| IG_VC0759 | 2 |
| VC0759 | 2 |
| VC0758 | 3 |
| VC0756 | 2 |
| VC0753 | 3 |
| VC0752 | 2 |
| VC0751 | 2 |
| IG_VC0750 | 2 |
| VC0750 | 2 |
| IG_VC0749 | 2 |
| VC0749 | 2 |
| IG_VC0748 | 2 |
| VC0748 | 2 |
| IG_VC0747 | 2 |
| VC0747 | 3 |
| VC0745 | 3 |
| IG_VC0733 | 3 |
| VC0733 | 3 |
| VC0727 | 3 |
| IG_VC0726 | 2 |
| VC0726 | 2 |
| VC0725 | 2 |
| VC0724 | 2 |
| VC0708 | 2 |
| VC0705 | 2 |
| IG_VC0704 | 3 |
| VC0696 | 2 |
| IG_VC0694 | 3 |
| VC0685 | 2 |
| VC0683 | 2 |
| IG_VC0682 | 2 |
| VC0682 | 2 |
| IG_VC0681 | 2 |
| VC0681 | 2 |
| VC0680 | 2 |
| IG_VC0679 | 3 |
| VC0679 | 2 |
| IG_VC0678 | 2 |
| VC0675 | 2 |
| IG_VC0674 | 2 |
| VC0674 | 2 |
| IG_VC0673 | 2 |
| IG_VC0664 | 3 |
| VC0664 | 2 |
| IG_VC0662 | 2 |
| IG_VC0661 | 3 |
| VC0660 | 3 |
| IG_VC0647 | 3 |
| VC0647 | 2 |
| IG_VC0646 | 3 |
| VC0646 | 2 |
| IG_VC0645 | 2 |
| VC0644 | 3 |
| VC0643 | 2 |
| VC0642 | 2 |
| VC0641 | 2 |
| VCt047 | 2 |
| VC0640 | 2 |
| VC0639 | 2 |
| IG_VC0638 | 2 |
| VC0638 | 2 |
| IG_VC0637 | 2 |
| VC0637 | 2 |
| IG_VC0636 | 3 |
| IG_VC0633 | 3 |
| IG_VC0631 | 2 |
| VC0631 | 2 |
| VC0627 | 2 |
| VC0626 | 2 |
| IG_VC0620 | 3 |
| VC0604 | 2 |
| IG_VC0603 | 3 |
| VC0594 | 2 |
| VC0593 | 2 |
| VC0592 | 2 |
| VC0591 | 2 |
| VC0581 | 2 |
| VC0576 | 2 |
| VC0573 | 3 |
| IG_VC0572 | 2 |
| VC0572 | 2 |
| VC0571 | 2 |
| VC0570 | 2 |
| IG_VC0569 | 2 |
| VC0569 | 2 |
| VC0565 | 2 |
| IG_VC0564 | 2 |
| VC0564 | 2 |
| IG_VC0563 | 2 |
| VC0563 | 2 |
| IG_VC0562 | 2 |
| VC0562 | 2 |
| IG_VC0561 | 2 |
| VC0561 | 2 |
| IG_VC0560 | 2 |
| VC0560 | 2 |
| IG_VC0559 | 3 |
| VCt041 | 2 |
| IG_VCt040 | 2 |
| VCt040 | 2 |
| IG_VCt039 | 2 |
| VCt039 | 2 |
| IG_VC0548 | 2 |
| VC0548 | 2 |
| IG_VC0547 | 3 |
| VC0545 | 2 |
| VC0541 | 2 |
| VC0540 | 3 |
| VC0539 | 3 |
| IG_VC0529 | 3 |
| VC0529 | 2 |
| VC0528 | 2 |
| VC0527 | 2 |
| VC0524 | 2 |
| IG_VC0520 | 3 |
| VC0520 | 2 |
| IG_VC0519 | 2 |
| VC0519 | 2 |
| IG_VC0517 | 2 |
| VCt038 | 2 |
| IG_VC0515 | 2 |
| VC0515 | 2 |
| IG_VC0514 | 2 |
| VC0514 | 3 |
| VC0513 | 3 |
| IG_VC0512 | 3 |
| VC0490 | 3 |
| IG_VC0489 | 3 |
| VC0487 | 2 |
| IG_VC0486 | 3 |
| VC0478 | 2 |
| IG_VC0477 | 3 |
| VC0477 | 2 |
| IG_VC0476 | 2 |
| VC0472 | 2 |
| IG_VC0466 | 2 |
| VC0466 | 2 |
| IG_VC0460 | 2 |
| VC0460 | 2 |
| VC0446 | 2 |
| IG_VC0445 | 2 |
| VC0445 | 2 |
| VC0444 | 2 |
| VC0443 | 2 |
| VC0440 | 2 |
| VC0437 | 2 |
| VC0436 | 2 |
| IG_VC0435 | 2 |
| VC0435 | 2 |
| IG_VC0434 | 3 |
| VC0434 | 2 |
| IG_VC0423 | 3 |
| VC0417 | 2 |
| VC0416 | 2 |
| IG_VC0415 | 2 |
| VC0415 | 2 |
| IG_VC0414 | 3 |
| VC0397 | 2 |
| IG_VC0396 | 3 |
| VC0386 | 2 |
| VC0385 | 2 |
| VC0384 | 2 |
| IG_VC0383 | 2 |
| VC0374 | 3 |
| VC0372 | 2 |
| VC0371 | 2 |
| VC0368 | 2 |
| VC0367 | 2 |
| VC0366 | 2 |
| IG_VC0365 | 3 |
| IG_VC0361 | 2 |
| VC0361 | 2 |
| IG_VC0360 | 2 |
| VC0360 | 2 |
| IG_VC0359 | 2 |
| VC0359 | 2 |
| IG_VC0358 | 2 |
| VC0358 | 2 |
| VC0357 | 2 |
| VC0356 | 2 |
| VC0344 | 3 |
| VC0343 | 2 |
| VC0341 | 2 |
| IG_VC0340 | 3 |
| VC0340 | 2 |
| VC0339 | 2 |
| IG_VC0336 | 3 |
| VC0336 | 2 |
| VC0332 | 2 |
| VC0329 | 2 |
| IG_VC0328 | 2 |
| VC0328 | 2 |
| IG_VC0327 | 2 |
| VC0327 | 2 |
| IG_VC0326 | 2 |
| VC0326 | 2 |
| IG_VC0325 | 2 |
| VC0325 | 2 |
| VC0324 | 2 |
| IG_VC0323 | 2 |
| VC0323 | 2 |
| IG_VC0322 | 2 |
| VC0322 | 2 |
| IG_VC0321 | 2 |
| VC0320 | 2 |
| VC0319 | 2 |
| VC0318 | 2 |
| VC0315 | 2 |
| VC0309 | 2 |
| VC0307 | 2 |
| VC0306 | 3 |
| IG_VC0297 | 3 |
| VC0297 | 2 |
| VC0296 | 2 |
| VC0295 | 2 |
| IG_VC0276 | 2 |
| VC0276 | 2 |
| VC0275 | 3 |
| VC0259 | 3 |
| IG_VC0258 | 3 |
| VC0258 | 3 |
| VC0249 | 3 |
| VC0248 | 2 |
| IG_VC0247 | 2 |
| VC0247 | 2 |
| VC0246 | 3 |
| VC0238 | 3 |
| VC0237 | 3 |
| VC0236 | 3 |
| VC0233 | 2 |
| VC0228 | 3 |
| VC0227 | 2 |
| VC0226 | 2 |
| VC0225 | 2 |
| VC0222 | 2 |
| VC0219 | 2 |
| VC0218 | 2 |
| IG_VC0217 | 3 |
| VC0215 | 2 |
| VC0213 | 2 |
| VC0211 | 3 |
| IG_VC0175 | 3 |
| IG_VC0168 | 3 |
| VC0167 | 2 |
| VC0162 | 2 |
| VC2755 | 2 |
| VC2762 | 2 |
| VC2763 | 2 |
| VC2764 | 2 |
| IG_VC2765 | 2 |
| VC2765 | 2 |
| IG_VC2766 | 2 |
| VC2766 | 2 |
| VC2767 | 2 |
| VC2768 | 2 |
| IG_VC2769 | 2 |
| VC2769 | 2 |
| IG_VC2770 | 2 |
| VC2770 | 2 |
| VC2771 | 2 |
| VC2775 | 2 |
| IG_VCA0001 | 2 |
| VCA0001 | 3 |
| IG_VCA0002 | 2 |
| VCA0002 | 2 |
| VCA0016 | 2 |
| VCA0021 | 2 |
| VCA0040 | 2 |
| IG_VCA0061 | 2 |
| VCA0061 | 2 |
| VCA0079 | 3 |
| VCA0093 | 3 |
| IG_VCA0094 | 2 |
| VCA0094 | 2 |
| IG_VCA0095 | 2 |
| VCA0095 | 3 |
| IG_VCA0105 | 3 |
| VCA0106 | 3 |
| VCA0124 | 2 |
| IG_VCA0125 | 2 |
| IG_VCA0143 | 2 |
| VCA0143 | 2 |
| IG_VCA0144 | 3 |
| IG_VCA0148 | 2 |
| VCA0162 | 2 |
| VCA0198 | 2 |
| VCA0199 | 3 |
| VCA0200 | 3 |
| VCA0201 | 3 |
| IG_VCA0202 | 3 |
| VCA0207 | 2 |
| IG_VCA0208 | 2 |
| VCA0208 | 2 |
| IG_VCA0219 | 3 |
| IG_VCA0247 | 2 |
| IG_VCA0287 | 3 |
| VCA0287 | 2 |
| IG_VCA0289 | 2 |
| VCA0289 | 2 |
| IG_VCA0290 | 2 |
| VCA0290 | 2 |
| IG_Intl4-VCA0291 | 3 |
| VCA0324 | 3 |
| IG_VCA0325 | 3 |
| VCA0333 | 2 |
| IG_VCA0334 | 2 |
| VCA0334 | 2 |
| VCA0360 | 2 |
| VCA0363 | 2 |
| IG_VCA0392 | 2 |
| VCA0392 | 2 |
| IG_VCA0477 | 3 |
| VCA0477 | 2 |
| VCA0486 | 2 |
| VCA0488 | 2 |
| VCA0498 | 3 |
| IG_VCA0524 | 3 |
| VCA0524 | 3 |
| VCA0528 | 2 |
| IG_VCA0570 | 3 |
| VCA0572 | 2 |
| IG_VCA0576 | 3 |
| VCA0614 | 3 |
| VCA0616 | 2 |
| VCA0628 | 2 |
| IG_VCA0643 | 2 |
| VCA0643 | 3 |
| IG_VCA0669 | 3 |
| VCA0669 | 3 |
| IG_VCA0693 | 3 |
| IG_VCA0701 | 2 |
| VCA0701 | 2 |
| IG_VCA0702 | 3 |
| IG_VCA0786 | 3 |
| VCA0786 | 2 |
| IG_VCA0812 | 3 |
| IG_VCA0834 | 2 |
| VCA0834 | 3 |
| IG_VCA0874 | 3 |
| VCA0874 | 3 |
| VCA0921 | 2 |
| VCA0925 | 2 |
| VCA0934 | 3 |
| IG_VCA0935 | 3 |
| IG_VCA0952 | 3 |
| VCA0952 | 2 |
| IG_VCA0953 | 3 |
| IG_VCA0975 | 3 |
| IG_VCA1037 | 3 |
| VCA1044 | 3 |
| IG_VCA1064 | 2 |
| VCA1064 | 3 |
| VCA1079 | 2 |
| IG_VCA1114 | 2 |
| VCA1114 | 2 |
| VCA1115 | 2 |
| IG_VCA0001 | 2 |

